# Image-Guided Targeting of Mitochondrial Metabolism Sensitizes Pediatric Malignant Rhabdoid Tumors to Low Dose Radiotherapy

**DOI:** 10.1101/2024.08.09.607364

**Authors:** Wenxi Xia, Matthew Goff, Carmine Schiavone, Neetu Singh, Jiemin Huang, Esther Need, Joseph Cave, David L. Gillespie, Randy L. Jensen, Mark D. Pagel, Prashant Dogra, Sixiang Shi, Shreya Goel

**Affiliations:** Department of Molecular Pharmaceutics, University of Utah, Salt Lake City, UT 84112, United States; Mathematics in Medicine Program, Department of Medicine, Houston Methodist Research Institute, Houston, TX, USA; Department of Chemical, Materials and Industrial Production Engineering, University of Naples Federico II, Naples, Italy; Physiology, Biophysics, and Systems Biology Program, Graduate School of Medical Sciences, Weill Cornell Medicine, New York, NY, USA; Department of Neurosurgery, Huntsman Cancer Institute, University of Utah, Salt Lake City, UT 84132, United States; Department of Medical Physics, University of Wisconsin-Madison, Madison, WI 53705, United States; Department of Physiology and Biophysics, Weill Cornell Medical College, New York, NY, USA; Department of Radiology and Imaging Sciences, University of Utah, Salt Lake City, UT 84112, United States; Department of Biomedical Engineering, University of Utah, Salt Lake City, UT 84112, United States

## Abstract

Tumor hypoxia leads to radioresistance and markedly worse clinical outcomes for pediatric malignant rhabdoid tumors (MRT). Our transcriptomics and bioenergetic profiling data reveal that mitochondrial oxidative phosphorylation (OXPHOS) is a metabolic vulnerability of MRT and can be exploited to overcome consumptive hypoxia by repurposing an FDA-approved anti-malarial drug, Atovaquone (AVO). We then establish the utility of Oxygen-Enhanced-Multispectral Optoacoustic Tomography (OE-MSOT), a label-free, ionizing radiation-free imaging modality, to visualize and quantify spatiotemporal changes in tumor hypoxia in response to AVO. We show a potent but transient increase in tumor oxygenation upon AVO treatment which results in complete elimination of tumors in all tested mice when combined with 10 Gy radiotherapy, a dose several times lower than the current clinic standard. Finally, we use translational mathematical modeling for systematic evaluation of dosing regimens, administration timing, and therapeutic synergy in a virtual clinical patient population. Together, our work establishes a framework for safe and pediatric patient-friendly image-guided metabolic radiosensitization of rhabdoid tumors.

## Introduction

Malignant rhabdoid tumor (MRT) and atypical rhabdoid tumor (rhabdoid tumors occurring in the brain) are rare but one of the most lethal soft tissue cancers with a median overall survival ranging from 6 to 17 months (*1*), affecting infants and young children (*2*) with approximately 20 to 25 new cases diagnosed each year in the U.S. (*3*). Given the rarity of these malignancies, new drug discovery is scientifically and financially challenging. Three conventional treatments are commonly employed for MRT, varying in part based on the disease stage: surgery, X radiation therapy (RT) and chemotherapy (*4*). Experimental treatment modalities including immunotherapy (*5, 6*) and targeted therapy (*7, 8*) are also being tested in preclinical trials (*9*). However, as most MRT patients are younger than three years old, special considerations are associated with current treatments. For example, chemotherapy is initiated after surgery but not desirable since it is often associated with heterogenous response and significant immediate and long-term toxicity in infant patients (*10*). All patients are treated with high dose localized RT. However, it is also associated with late-stage side-effects such as growth and developmental delays in pediatric patients (*11*), as well as gastrointestinal dysfunction, pulmonary issues, infertility, cardiac abnormalities, and the potential for secondary cancers later in life (*12*). Thus, a safe and effective radiosensitizing adjuvant treatment that maximizes RT efficiency at lower doses and minimizes radiotoxicity can become a “game-changer” in pediatric MRT management.

Nearly all solid tumors suffer from diminished oxygen availability or tumor hypoxia (*13*), resulting from increased oxygen consumption because of high metabolic demand and inefficient oxygen delivery due to leaky and compressed vessels (*14*). Tumor hypoxia appears to be strongly associated with cancer progression, including epithelial-mesenchymal transition, tumor vascularization, angiogenesis, cell migration/invasion, metastasis, immunosuppression (*15, 16*), as well as poor outcomes for various treatment modalities including RT, as oxygen acts as a potent radiosensitizer, enhancing the DNA damage caused by ionizing radiation (*17, 18*). Therefore, attenuation of hypoxia may be a highly appealing strategy to enhance the sensitivity of tumors to radiation and thereby improve therapeutic outcomes in pediatric MRT patients.

Currently, there are two common strategies employed to relieve tumor hypoxia: increasing oxygen delivery by exogenous routes or vascular remodeling, or, reducing demand for oxygen by modulating cellular oxygen consumption rate (OCR) through metabolic reprogramming (*19–21*).. Our transcriptomic profiling of MRT patient data in the TARGET database revealed a significant upregulation of the oxidative phosphorylation (OXPHOS) pathways, suggesting that reducing hypoxia through OXPHOS inhibition could be a clinically feasible approach to enhance radiosensitivity in MRT treatment. Atovaquone (AVO), the first US Food and Drug Administration (FDA)-approved anti-malarial and anti-pneumocystis pneumonia drug, was recently repurposed as an OXPHOS inhibitor for tumor growth suppression (*22*). AVO reduces OCR by inhibiting the mitochondrial electron transport chain at complex III, and studies have reported that AVO could effectively alleviate tumor hypoxia in various adult tumor types, as a monotherapy, as well as combination with radiotherapy and immunotherapy (*23, 24*). We reasoned that AVO could be the game-changing radiosensitizer to enable effective treatment of pediatric solid tumors at significantly lower radiation doses, which has not been tested yet.

To investigate the effectiveness of AVO in mouse models of pediatric MRT and characterize the extent and duration of hypoxia modulation, we employed multispectral optoacoustic tomography (MSOT), a multiscale imaging modality to noninvasively visualize oxygenation and vascular perfusion at the tumor site (*25–27*). Data obtained at various wavelengths in optoacoustic tomography enables the quantification of oxy- and deoxy-hemoglobin levels which can be employed to measure oxygen saturation (%sO_2_) of the blood (*28, 29*). By performing %sO_2_ measurement following a switch of respiration gas from air (21% oxygen) to 100% oxygen, using the oxygen-enhanced (OE-MSOT) procedure, ΔsO_2_ can be measured, that represents the “available oxygen capacity” in the tumor (*30*). Additionally, dynamic contrast enhanced (DCE)-MSOT can also be performed at the same time to evaluate tumor perfusion by analyzing NK^trans^ and K_ep_ pharmacokinetics (PK) parameters. These parameters respectively reflect the wash-in and wash-out perfusion rates of an FDA-approved agent, indocyanine green (ICG), within the tumor (*31*). Thus, OE-and DCE-MSOT together can provide quantitative, multiparametric, real-time functional information about tumor responses to administered AVO and RT in live animals.

In this work, we integrate whole body in vivo optoacoustic imaging and statistical parametric mapping with tissue- and cellular scale histological analysis to predict and assess the effectiveness of AVO-potentiated low dose RT therapy in A-204 MRT xenografts **(Scheme 1)**. Our findings underscore the efficacy of OE-MSOT in mapping the spatiotemporal changes of drug pharmacodynamics, encouraging its use to guide personalized combination therapy regimens. Notably, our results indicate that at low pharmacologically achievable concentrations, AVO can effectively but transiently alleviate hypoxia in A-204 tumors, and strongly inhibit tumor growth in combination with a single 10 Gy X ray dose with no adverse effects. These insights provide an important contribution towards pediatric MRT management in the clinic for which limited therapies exist. Further, to bridge the gap between preclinical findings and clinically applicable treatment regimens, mathematical modeling is employed as a complementary approach to experimental investigations. By simulating tumor dynamics in virtual patient populations, the framework enabled a systematic evaluation of dosing regimens, administration timing, and therapeutic synergy. This approach identified optimal combination protocols to enhance clinical outcomes, guiding future studies and potentially supporting the translation of AVO as a radiosensitizer in MRT.

**Scheme 1:**
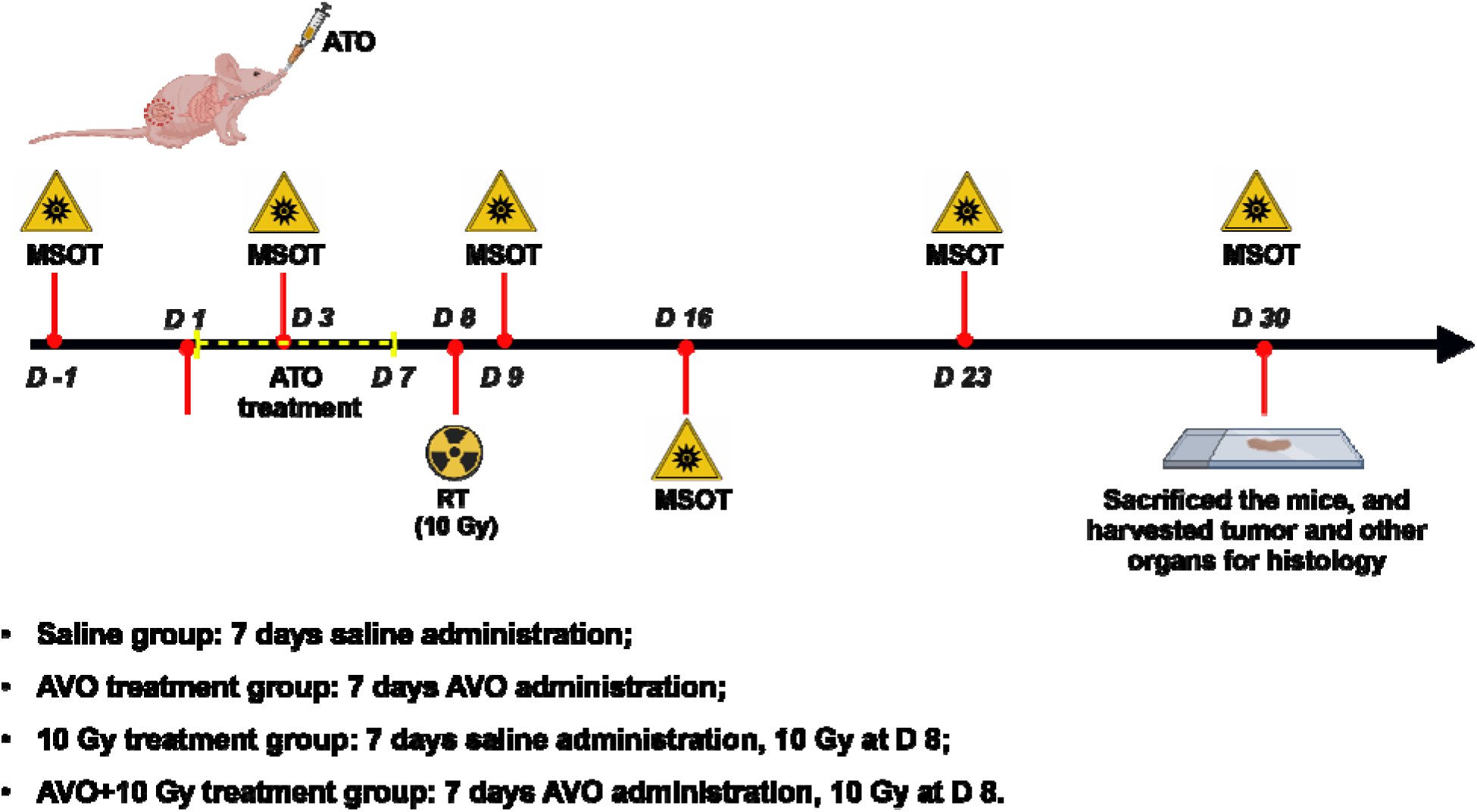
Treatment and imaging plan of A-204 MRT xenografts. Once the tumor volume reached 50 mm^3^, MSOT scans were performed before (Day-1), during (Day 3), and after (Day 9, 16, 23 and 30) initiating treatment for all groups. Mice were euthanized when the tumors reached 1000 mm^3^, or when moribund or at day 60 post-treatment. Tumors and other organs were harvested for subsequent histology analysis. Created with BioRender.com.

## Results

### OXPHOS pathway is upregulated in clinical rhabdoid tumor patients

Tumor hypoxia plays a key role in increasing resistance to RT, leading to significantly worse clinical outcomes in pediatric MRT treatment. Thus, our objective here is to investigate targetable pathways that can reduce tumor hypoxia, enhance the effectiveness of RT, and minimize long-term radiotoxicity. Towards this end, we profiled transcriptomics data from 63 rhabdoid cancer patients from the GDC database. The workflow for the RNA sequencing, pathway enrichment and survival curve plotting were shown in **Fig. 1A**. After conducting ssGSEA on the transcriptomic data using Hallmark gene sets, we discovered that the OXPHOS pathway exhibited higher enrichment level compared to other pathways in rhabdoid tumor patients, regardless of cancer stage (**Fig. 1B**). Therefore, we hypothesized that inhibiting OXPHOS could serve as an actionable strategy to augment RT by attenuating consumptive hypoxia. Additionally, to further understand which genes in the OXPHOS pathway influence survival rates, we performed Cox proportional hazards regression modeling on all OXPHOS related genes and identified 11 genes with significant p-values (p < 0.05, **Fig. 1C**), including genes associated with complex I and III activity in the electron transport chain (ETC). As shown in **Fig. 1D** and **S1**, survival curve analysis revealed that patients with high expression of UQCRFS1 (component of ubiquinol-cytochrome c oxidoreductase involved in complex III assembly) and NDUFA3 (involved in complex I assembly) had significantly lower survival rates compared to those with lower expression. Disrupting the complexes along the ETC is the most clinically viable way to pharmacologically target OXPHOS and many OXPHOS inhibitors targeting complexes I and III are in various stages of clinical trials. Atovaquone (AVO), a competitive inhibitor of ubiquinone, a crucial component of complex III (cytochrome bc1 complex (*32*)) was selected in this study (**Fig. 1E**). AVO is an FDA-approved anti-parasitic drug with a well-established safety and efficacy profiles in both adults and children (*33*). Importantly, AVO has been shown to inhibit OXPHOS at pharmacologically achievable concentrations in adult tumors (*34*).

**Fig. 1.**
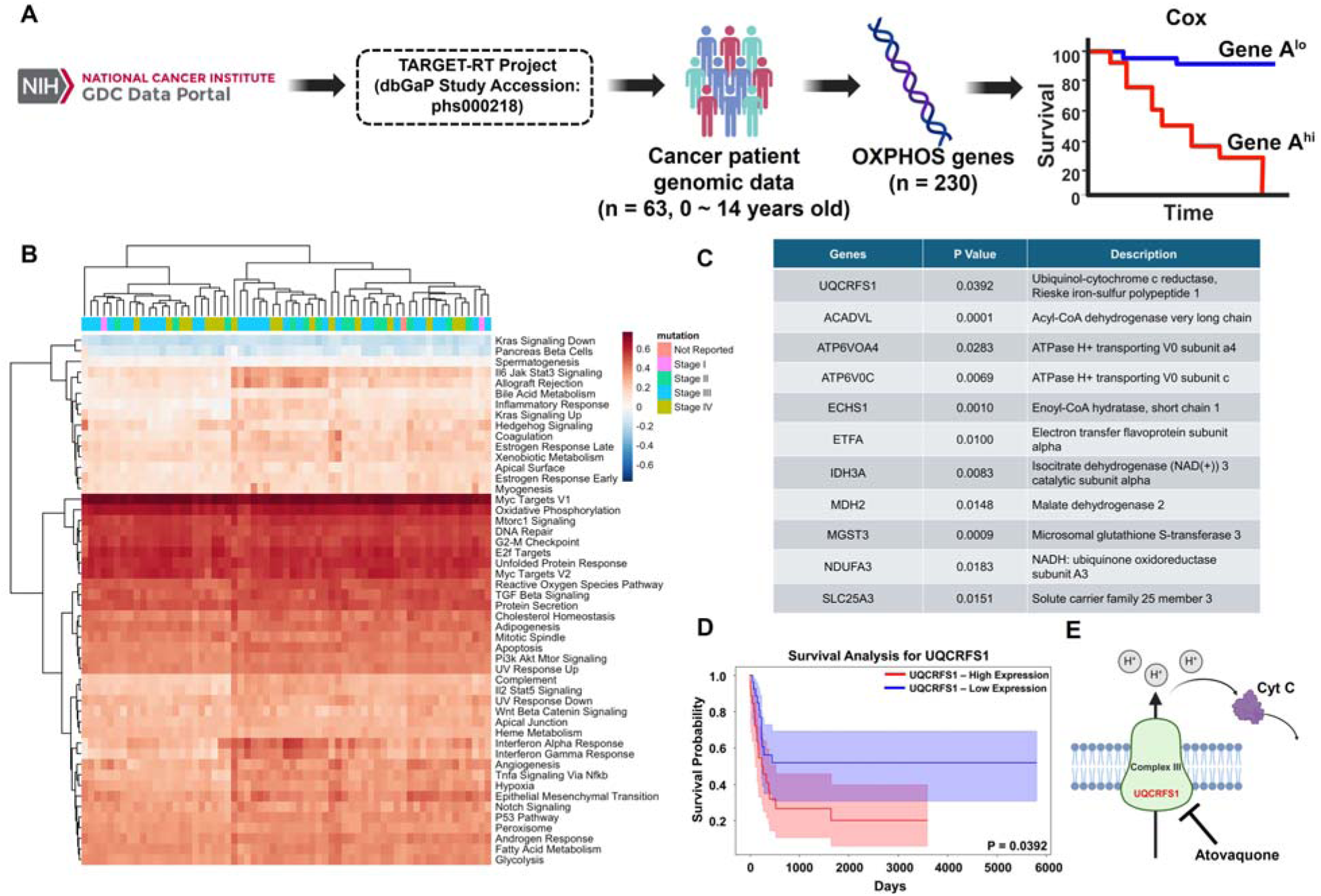
**(A)** The scheme of the RNA sequencing, pathway enrichment and survival curve plotting. **(B)** The heatmap indicated a significant upregulation of the OXPHOS pathway across different stages of rhabdoid tumor patients. **(C)** P values and description of selected 11 OXPHOS-related genes. **(D)** Kaplan-Meier survival analysis showed that patients with high UQCRFS1 expression had a significantly worse survival rate compared to those with low expression. **(E)** Schematic showing the UQCRFS1 gene is associated with mitochondrial complex III, inhibited by AVO.

### AVO effectively alleviates hypoxia by inhibiting mitochondrial respiration in A-204 cells

To verify the inhibitory effect of AVO on mitochondrial respiration in pediatric MRT, we measured the oxygen consumption rates (OCR) in A-204 cells using a Seahorse XF Pro analyzer. Control groups treated with DMSO alone were processed in parallel. Before measuring OCR traces, cytotoxicity experiments were conducted to determine the approximate AVO concentration for Seahorse experiments. As shown in **Fig. 2A**, AVO monotherapy did not exert substantial cytotoxicity on A-204 cells, yielding an EC_50_ value ∼28.36 μM. However, this concentration was sufficient to significantly suppress OXPHOS. Through the sequential injection of oligomycin, FCCP, and rotenone/antimycin A, the basal respiration, maximal respiration and ATP production of A-204 cells were recorded, respectively. OCR traces shown in **Fig. 2B-E** demonstrated that AVO-treated groups exhibited significant suppression in basal respiration, maximal respiration and ATP production compared with the control groups, indicating AVO can effectively inhibit mitochondrial respiration. The metabolic impact of AVO treatment was assessed by plotting ECAR and OCR to evaluate shifts in the metabolic state of A-204 cells. AVO treatment elicited a distinct metabolic response. Specifically, A-204 cells transitioned from an energetic state to a quiescent state (**Fig. S2**). Notably, this transition occurred without a significant increase in glycolysis, as evidenced by stable ECAR values and the lack of a shift toward the glycolytic quadrant in the metabolic profile. These results suggest that AVO treatment avoids the excessive glycolytic activation associated with other OXPHOS inhibitors, reducing the risk of lactic acidosis and offering a safer metabolic profile.

**Fig. 2.**
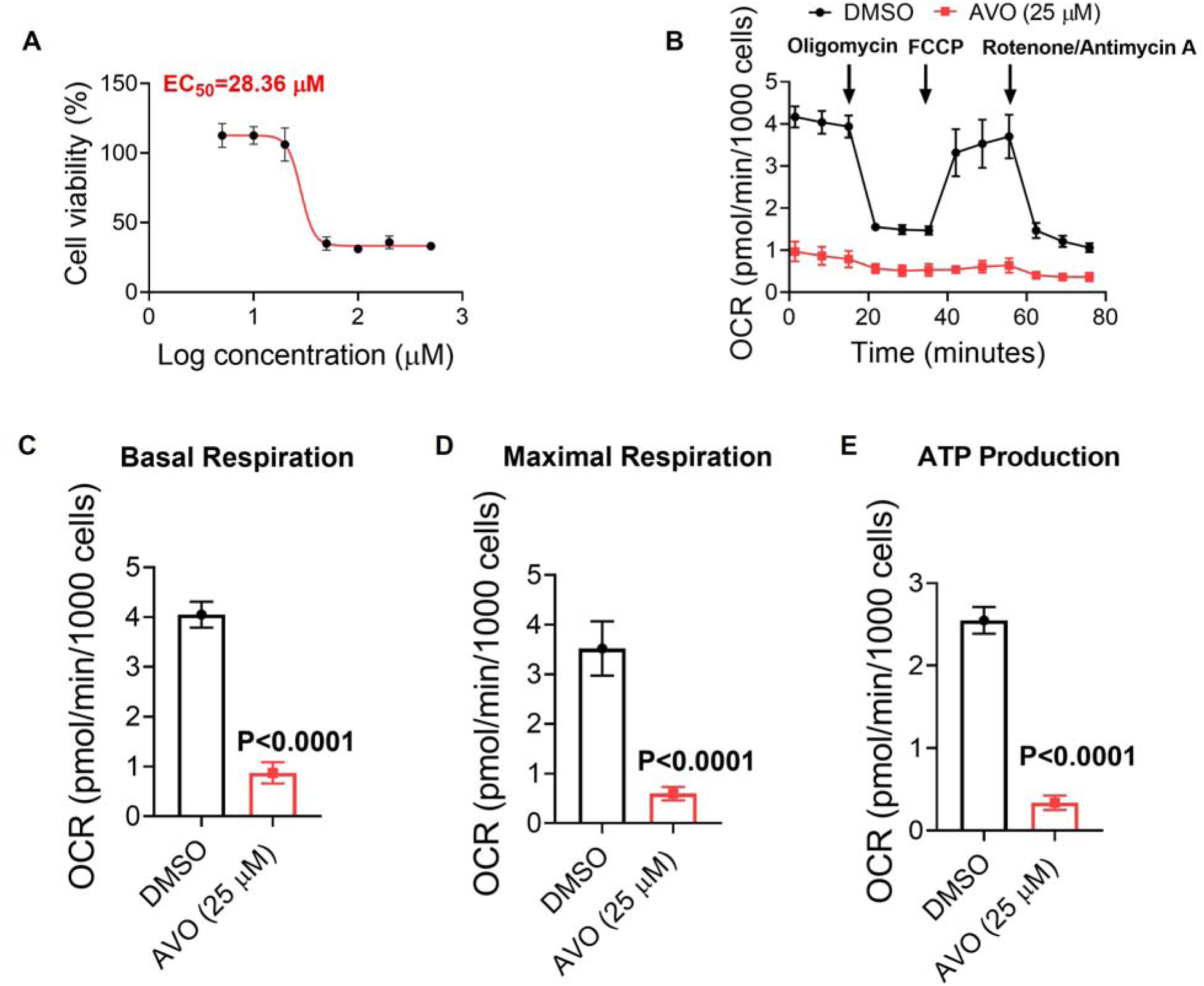
AVO inhibits mitochondrial respiration of A-204 cells. **(A)** Dose response curves depicting the effect of AVO on viability of A-204 cells as tested by MTT assay. Seahorse experiment indicating **(B)** OCR traces of A-204 cells treated with vehicle (DMSO) or AVO. **(C, D, E)** Significant reductions in basal respiration, maximal respiration, and ATP production were observed in AVO treated groups when compared to vehicle-treated controls. All OCR measurements were normalized by Hoechst 33342 staining. Data are presented as mean ± SD.

We also compared the performance of AVO with metformin, a commonly applied OXPHOS inhibitor. As seen in **Fig. S3**, the EC_50_ value of metformin for A-204 cells was found to be 13.15 mM, which was significantly higher than that of AVO (∼28.36 µM). This confirms that AVO could potentially lead to better therapeutic effects and less off-target toxicity at lower pharmacological concentrations, suggesting improved *in vivo* performance when compared to metformin.

### *In vivo* OE-MSOT shows that AVO treatment improves tumor oxygenation transiently

We next investigated the *in vivo* pharmacodynamics of AVO treatment on A-204 xenografts using longitudinal OE-MSOT. OE-MSOT maps tumor oxygenation based on changes in hemoglobin oxygen saturation (%SO_2_, computed as a ratio of HbO_2_ to total Hemoglobin) following a switch in delivery from 21% O_2_ (medical air) to supplemental (100%) O_2_. The ΔsO_2_ parameter is computed as the difference between %sO_2_^O2^ and %sO_2_^air^. Compared to static measurements, the dynamic nature of OE-MSOT has been found to provide more robust and reproducible spatiotemporal evaluations of tumor oxygenation (*35–37*). We hypothesized that hypoxic tumors will be refractory to hyperoxic gas breathing as high oxygen consumption will increase the oxygen extraction rate, resulting in a reduction of hemoglobin bound O_2_ (HbO_2_). Meanwhile, reduction in metabolic oxygen consumption will require a reduction in oxygen extraction, resulting in greater affinity of oxygen for hemoglobin, and hence higher %sO ^O2^ after hyperoxic gas challenge. Thus, blocking mitochondrial function by AVO should lead to reduced O_2_ consumption, and hence, higher ΔsO_2_.

Application of oxygen gas challenge on tumors at baseline (referred to as Day-1, when tumors reached ∼ 50 mm^3^) indicated negligible changes in the MSOT derived %sO_2_ indicating the hypoxic nature of the tumors (**Fig. S4**). **Fig. 3B** and **C** show that AVO treated tumors depicted a nearly 2-fold increase in ΔsO_2_ from baseline (ΔsO_2_ = 0.047 ± 0.02) to Day 3 (ΔsO_2_ = 0.094 ± 0.05) when compared to saline-treated controls (ΔsO_2_ = 0.050 ± 0.002 at Day -1 and 0.030 ± 0.011 at Day 3). A sharp decline in ΔsO_2_ can be seen from Day 3 to Day 9 in AVO-treated tumors, after AVO treatment was stopped at Day 7, indicating a gradual reversion to a hypoxic phenotype.

**Fig. 3.**
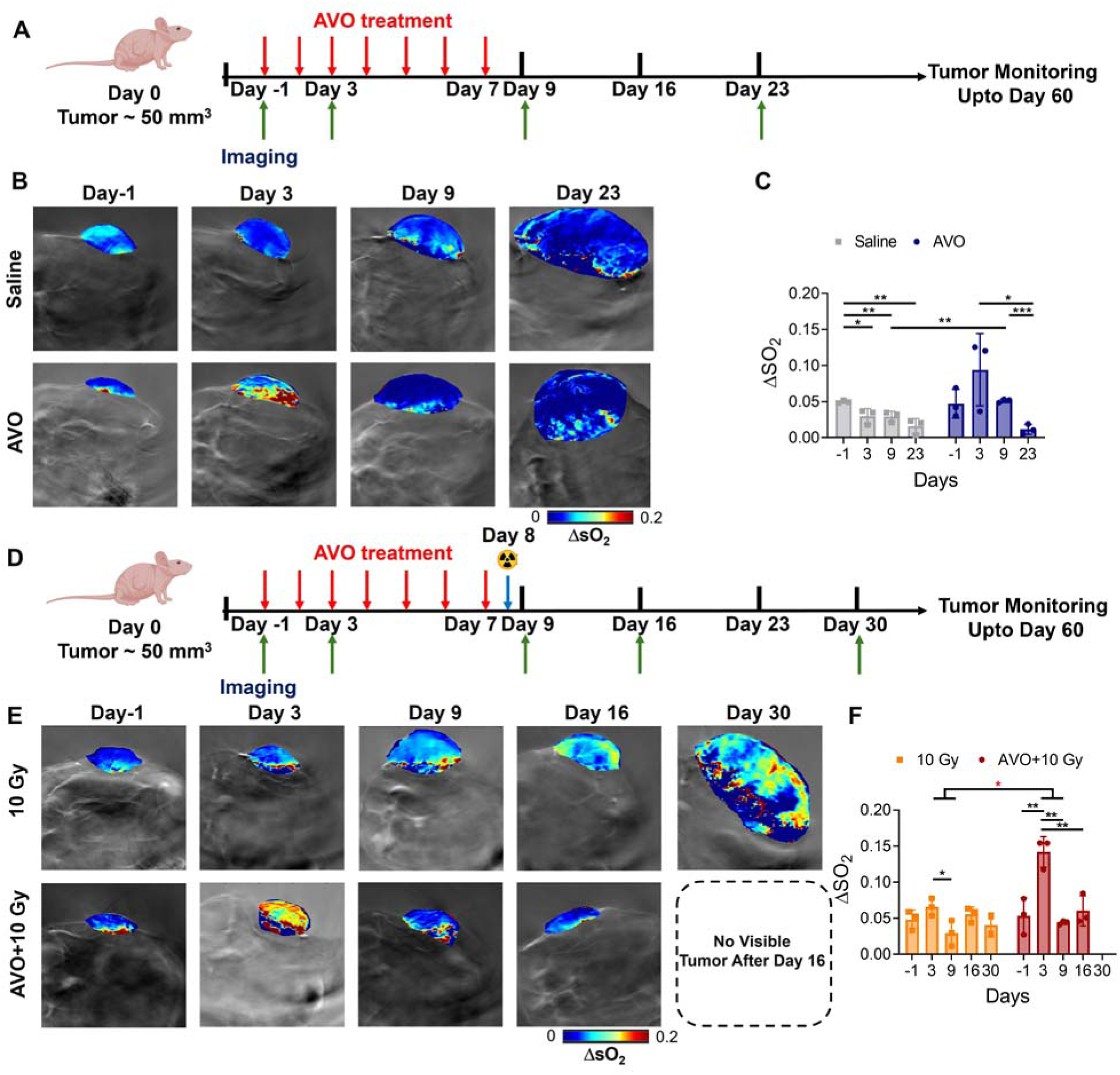
OE-MSOT shows that AVO treatment improves tumor oxygenation transiently on A-204 xenografts. **(A)** Treatment and imaging timeline for saline or AVO-treated groups. (n=3). **(B)** Parametric maps and **(C)** quantitative analysis of tumor ΔsO_2_ of saline and AVO treatment groups at each timepoint. **(D)** Treatment and imaging timeline for 10 Gy or AVO+10 Gy-treated groups. (n=3). **(E)** Parametric maps and **(F)** quantification analysis of tumor ΔsO_2_ of 10 Gy and AVO+10 Gy treatment groups at each time point. Data are presented as mean ± SD. Statistical analysis as indicated.

However, a statistically significant difference in ΔsO_2_ can still be observed between AVO and saline-treated mice at Day 9 (p=0.0074). OE-MSOT at endpoint (Day 23 for AVO treated group) showed a strong reduction in ΔsO_2_ (0.011 ± 0.007) which is comparable to saline-treated controls at endpoint (Day 30; 0.016 ± 0.011). This decrease in ΔsO_2_ can be related to continued tumor growth after cessation of therapy, at indicated by increase in tumor volume. Increased pimonidazole immunofluorescence (**Fig. 4A, 4B** and **Fig. S5**) validated our *in vivo* OE-MSOT observations.

**Fig. 4.**
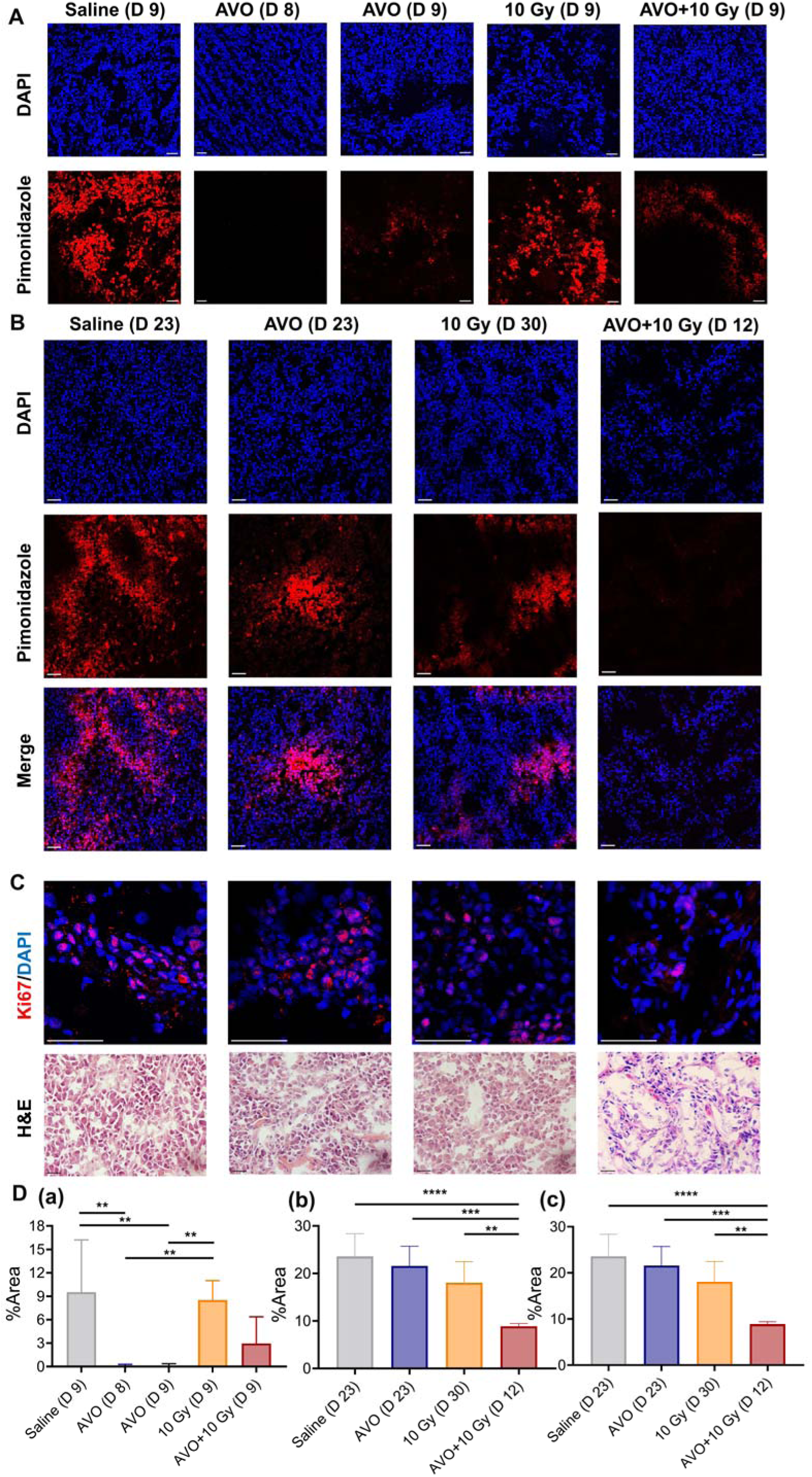
Immunofluorescence imaging shows that AVO alleviates tumor hypoxia. Immunofluorescence imaging of pimonidazole staining of tumors harvested at day 8 and 9 (mid-treatment) **(A)**, and endpoint **(B)** for different treatment groups. **(C)** Ki-67 and H&E staining at endpoint for different groups.. Scale bar = 50 µm. (**D)** Quantitative analysis of pimonidazole at (a: Day 9, b: end point) and Ki-67 at endpoint (c) in all treatment groups. Data are presented as mean ± SD.

Our results highlight the importance of longitudinal monitoring of hypoxia modulation by AVO using OE-MSOT. As the duration of hypoxia alleviation by AVO is short-lived and wanes off after the treatment is stopped, it is imperative that combination with radiotherapy is timed appropriately, so that the tumors are irradiated when they are most oxygenated. After establishing the oxygenation dynamics, a separate cohort of AVO-treated tumor-bearing mice were subjected to a single low 10 Gy dose of focused X-radiation, one day after AVO treatment was stopped. *Ex vivo* pimonidazole staining confirmed the well-oxygenated status of tumors at Day 8 (**Fig. 4A** and **Fig. 4D(a)**) Immediate (Day 9) and longer-term changes in tumor oxygenation following AVO+10 Gy combination treatment were assessed with OE-MSOT and immunofluorescence and compared to mice receiving 10 Gy irradiation only. As seen in **Fig. 3E** and **F**, irradiation of AVO-treated tumors results in an immediate and strong decrease in ΔsO_2_ (0.044 ± 0.002) at Day 9 compared to ΔsO_2_ (0.142 ± 0.021) at Day 3 (p value =0.0013), presumably caused by enhanced cellular death. We hypothesized that increased oxygenation by AVO bolstered greater ROS-mediated cell killing by low dose radiation. Interestingly, we observed a slight increase in tumor oxygenation a few days after irradiation, as measured by both OE-MSOT and pimonidazole immunostaining (**Fig. 4B**, **4D(b)** and **Fig. S5**). It is reasonable to assume that increased cellular apoptosis resulted in reduced oxygen consumption and hence a temporary increase in intratumoral oxygen levels. Ki-67 staining and H&E on Day 12 validated our premise (**Fig. 4C** and **Fig. 4D(c)**). To confirm that hypoxia modulation by AVO was tumor specific, and to confirm that pimonidazole staining truly reflected tissue oxygenation, we assessed major organs (heart, liver, kidney and muscle) in saline and AVO+10 Gy treated groups. As seen in **Fig. S6**, no significant difference was observed in pimonidazole staining for any of the tissues between saline and the combination treatment groups.

In contrast, tumors treated with irradiation only remained uniformly hypoxic with no significant changes in ΔsO_2_ values, consistent with resistance to low dose radiotherapy (**Fig. 3E** and **Fig. 3F**). Pimonidazole staining of irradiated tumors on Day 9 and endpoint indicated similar hypoxia levels as saline treated groups (**Fig. 4A** and **Fig. 4B**), which further corroborated our observations *in vivo*. Taken together, our data indicates the potential utility of monitoring oxygenation by OE-MSOT and ΔsO_2_ as a noninvasive biomarker to predict and evaluate tumor response to hypoxia modulating therapies and radiotherapy. As MSOT involves light and ultrasound for noninvasive interrogation of tissues, we believe it could be an attractive modality for imaging-based treatment planning and evaluation in children with cancers.

### AVO enhances the sensitivity of A-204 tumors to low dose radiation therapy

Given the significant improvement in tumor oxygenation after AVO treatment observed by OE-MSOT and pimonidazole staining, we hypothesized that AVO could enhance the sensitivity of A-204 tumors to low dose radiation therapy and improve tumor responses. Thus, after imaging, mice in monotherapy and combination therapy groups were monitored for tumor growth and survival over 60 days (**Fig. 5A**). Mouse body weight was also measured, as weight changes serve as a valuable parameter for assessing treatment toxicity *in vivo* (*38*). As shown in **Fig. 5B**, the body weight curves exhibited no significant difference among all treatment groups, suggesting that AVO, 10 Gy or combined treatments did not cause noticeable systemic toxic effects on mice, compared to saline treated controls. Tumor volume changes (**Fig. 5C**) showed that the average time required for tumors to reach 1000 mm^3^ was 29 days for saline-treated group, 30 days for AVO-treated group and 42 days for 10 Gy-treated group. One tumor in the RT only group had excellent response. However, for the AVO+10 Gy-treated group, all the tumors were completely abolished within 10 days post-treatment, and most strikingly, tumors did not recur for the remainder of the study. Endpoint tumor volume changes as measured by calipers (**Fig. 5D**) or observed visually (**Fig. S7** and **Fig. 5E**) demonstrated significant abrogation in all mice treated with AVO+10 Gy when compared to monotherapy or saline treated mice (p value <0.0001). As expected, the stellar tumor inhibitory effects of AVO + RT combination therapy, coupled with minimal toxicity resulted in significantly extended animal survival (**Fig. 5F**).

**Fig. 5.**
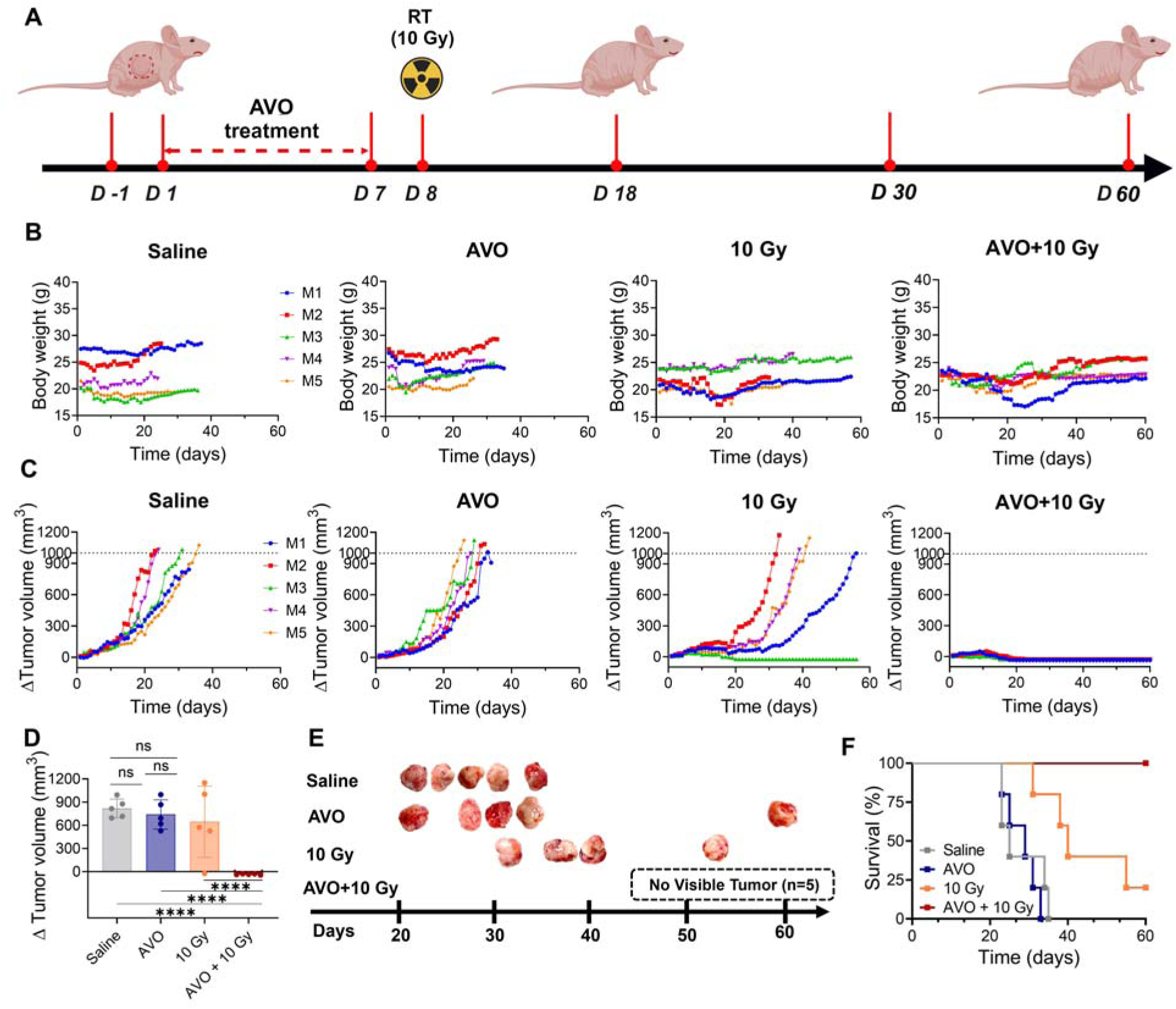
AVO increases low dose radiation treatment response in A-204 xenografts. **(A)** Treatment scheme of AVO+10 Gy combination therapy group. Graphs depicting **(B)** body weight and **(C)** tumor volume changes over 60 days in all treatment groups (n=5). ΔTumor volumes was computed by subtracting tumor volumes at baseline from the tumor volume at each time point. **(D)** ΔTumor volumes and (**E**) Photographs of excised tumors at endpoint for all groups. No visible tumors were found in AVO+10 Gy-treated groups on day 60. (**F**) Kaplan-Meier survival curves forall treatment groups. Data are presented as mean ± SD. Statistical analysis as indicated in the Methods section.

We used γ-H2AX as a biomarker of DNA damage to evaluate irradiation response in A-204 tumors (*39*). As seen in **Fig. 6A** and **B**, AVO+10 Gy-treated tumors depicted more obvious γ-H2AX signal when compared to saline-treated (p=0.0038), AVO-treated (p=0.0011) and 10 Gy-treated groups (p=0.0284; **Fig. 6C(b)**), demonstrating the radiosensitizing effect of AVO on typically radioresistant tumors. Subsequently, H&E staining of other organs including heart, liver, kidney and muscle was performed to further evaluate the treatment toxicity. As shown in **Fig. 6D**, no detectable necrosis or morphological changes were observed in these organs, which is consistent with the conclusions drawn from the body weight results.

**Fig. 6.**
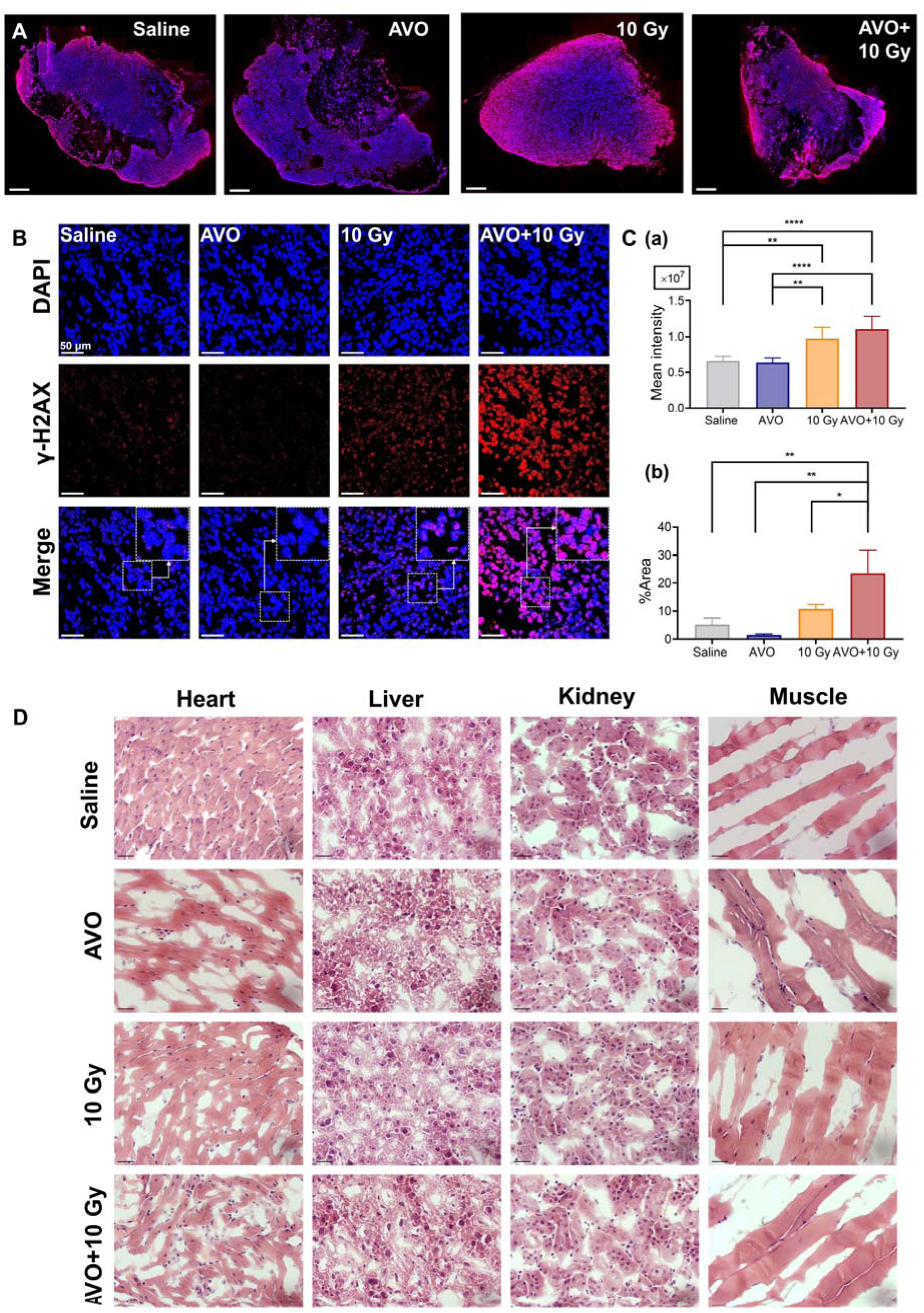
AVO enhanced DNA damage caused by RT. **(A)** Whole tissue imaging and (**B**) confocal micrographs of γ-H2AX immunofluorescence staining of all treatment groups on Day 9. **(Insets)** Stained A-204 tissue at high magnification. Scale bar = 50 µm for confocal images and scale bar = 1000 µm for whole tumor sections. **(C)** Quantification of γ-H2AX staining in whole tumor sections (**a**) and confocal micrographs (**b**) of all treatment groups. **(D)** H&E staining for heart, liver, kidney and muscle from various treatment groups. Scale bar=100 µm. Data represented as mean ± SD.

To test whether increased radiation doses would elicit tumor inhibitory effects, a separate cohort of A-204 xenografted mice were treated with 30 Gy dose (fractionated over 5 days, 6 Gy per fraction). Another group of mice was treated with AVO and fractionated RT dose (treatment schema shown in **Fig. S8**). As seen in **Fig. S8A**, a fractionated 30 Gy dose was well-tolerated (no apparent change in body weight) but only succeeded in slowing the tumor growth. In one mouse, the tumor did not respond to irradiation, as indicated by the exponential increase in tumor volume. On the hand, when combined with AVO, a 30 Gy fractionated dose exerted significant tumor control, with tumors disappearing within 5 days of stopping treatment (**Fig. S8B**).

Taken together, the data demonstrate that although AVO does not exert therapeutic effect as a monotherapy for MRT, the combination of AVO and low dose RT can be an effective and efficient strategy to maximize responses, with potential for clinical application in MRT treatment in the future. Our findings are significant for children with MRT through two avenues. First, we demonstrate that AVO can serve as a safe, FDA approved adjuvant to significantly enhance therapeutic efficacy of radiotherapy in otherwise radioresistant tumors and increase overall recurrence-free survival. Second, the ability to obliterate tumors at low radiation doses serves to greatly assuage concerns about longer-term radiotoxicities or secondary tumor growth, which are known to significantly impact quality of life and survival in patients subjected to aggressive chemoradiation therapies, earlier in childhood.

### DCE-MSOT indicates that AVO-mediated reduction in tumor hypoxia does not depend on vascular normalization

Prolonged hypoxia could result from irregular tumor vascularization in aggressive solid tumors (*40*), which compromises tumor perfusion and hinders effective oxygen delivery (*41*).To investigate whether AVO treatment could relieve tumoral hypoxia by vascular normalization, we employed DCE-MSOT to visualize and quantify changes in tumor vascular perfusion. **Fig. 7A** displays the parametric maps of normalized wash-in rate (NK^trans^) and wash-out rate (K_ep_) of ICG for the saline and AVO-treated groups on Day-1 (baseline), and Days 3, 9 and 23. There were no significant differences detected in NK^trans^ and K_ep_ values before, during, and after AVO treatment (**Fig. 7B**). Kinetic curves plotted for ICG accumulation over 15 minutes of dynamic scan demonstrated no apparent differences between tumors treated with saline or AVO (**Fig. 7C**). These results, together with our *in vitro* bioenergetics stress test, *in vivo* OE-MSOT and *ex vivo* histopathological data establish that improved tumor oxygenation by AVO is mediated by associated reduction in metabolic demand for oxygen and not through vascular normalization. Furthermore, we also compared the parametric maps, NK^trans^ and K_ep_ values, and ICG signal kinetics in the tumors subjected to 10 Gy monotherapy and AVO+10 Gy combination treatment groups. The results of **Fig. S9A** and **B** indicate that no meaningful difference was observed between the different groups. Interestingly, although changes in vascular perfusion are commonly observed after RT in solid tumors, we did not find any significant alterations in our study, possibly attributed to the intrinsic radioresistance of MRT to low dose radiation.

**Fig. 7.**
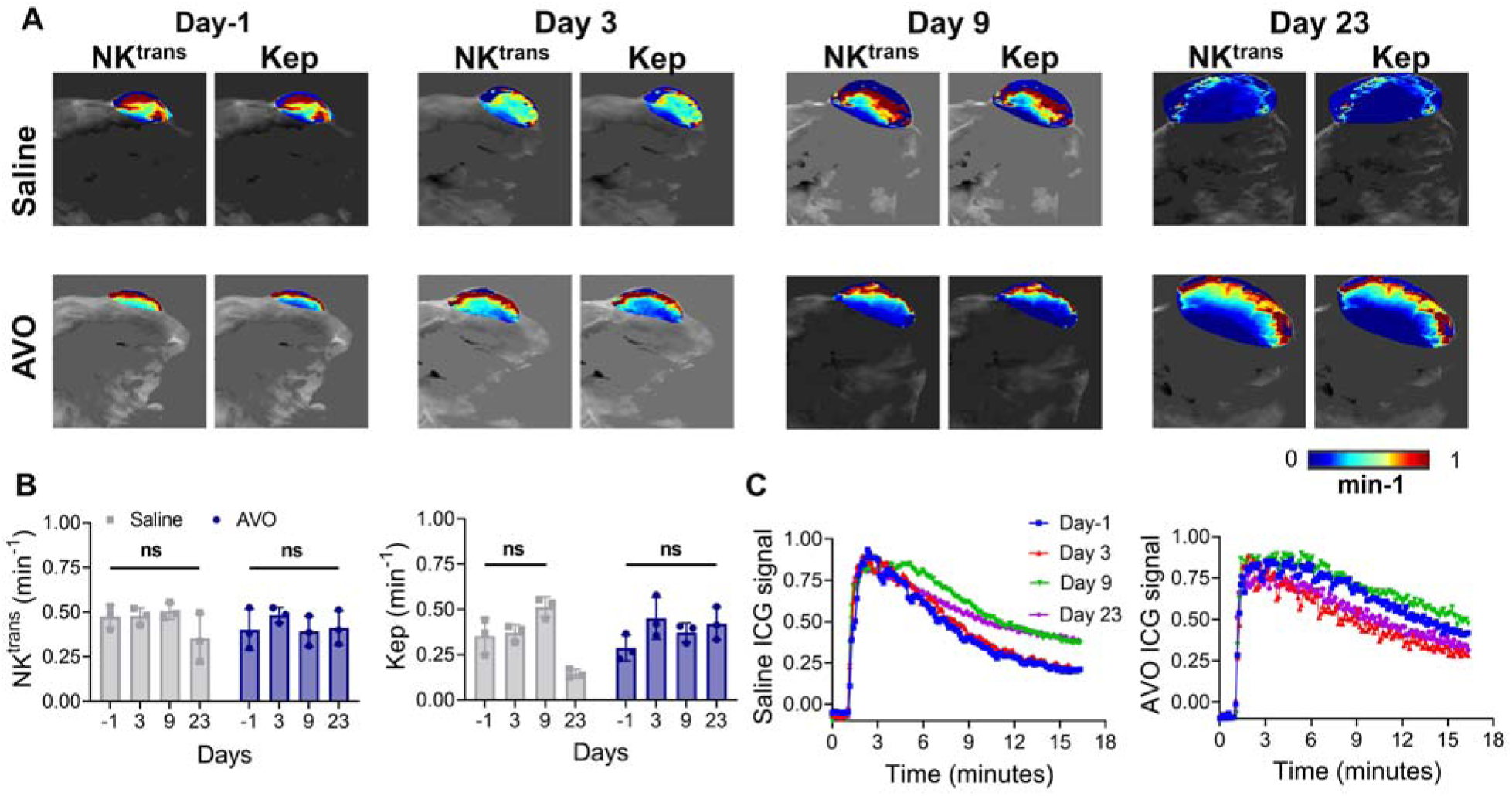
DCE-MSOT shows that AVO has no impact on tumor perfusion. **(A)** Parametric maps and **(B)** quantification analysis of NK^trans^ and K_ep_ for saline and AVO treated A-204 xenografts on Day-1, Day 3, Day 9 and Day 23. (n=3) **(C)** Kinetic curves depicting ICG accumulation in tumors over time in saline and AVO treatment groups. Data presented as mean ± SD.

Immunofluorescence imaging of CD31 was subsequently conducted on excised tumor tissues at Day 9 (**Fig. S9C**) and endpoint **(Fig. S10)** to validate our *in vivo* findings. Consistent with the DCE-MSOT results, CD31 staining did not show any significant difference among different treatment groups, confirming that AVO had no impact on vessel structure or functionality.

### OE-MSOT can distinguish between Atovaquone-sensitive and Atovaquone-resistant A-204 xenografts

To confirm that AVO was indeed essential for the observed therapeutic efficacy of low dose RT, we developed AVO-resistant A-204 cell line (A^R^-A-204) by treating cells to escalating doses of AVO over time (**Fig. 8A**). OCR and ECAR traces were first assessed in A^R^-A-204 cells to establish baseline. The results shown in **Fig. 8B** indicated that continuous AVO treatment resulted in lower baseline levels of mitochondrial respiration in A^R^-A-204 cells, significantly reducing basal respiration, maximal respiration, and ATP production (p value <0.0001, **Fig. 8D-F**) when compared to naïve A-204 cells. While ECAR traces showed no significant differences in aerobic glycolysis between A-204 and A^R^-A-204 cells (**Fig. 8C**), marked increase in glycolytic capacity (p value =0.0002) and glycolytic reserve (p value <0.0001, **Fig. S11A**) were observed alongside a metabolic shift to a quiescent state in A^R^-A-204 cells. Our results indicate metabolic plasticity associated with treatment resistance in MRT and warrant further examination into the metabolic effects of AVO.

**Fig. 8.**
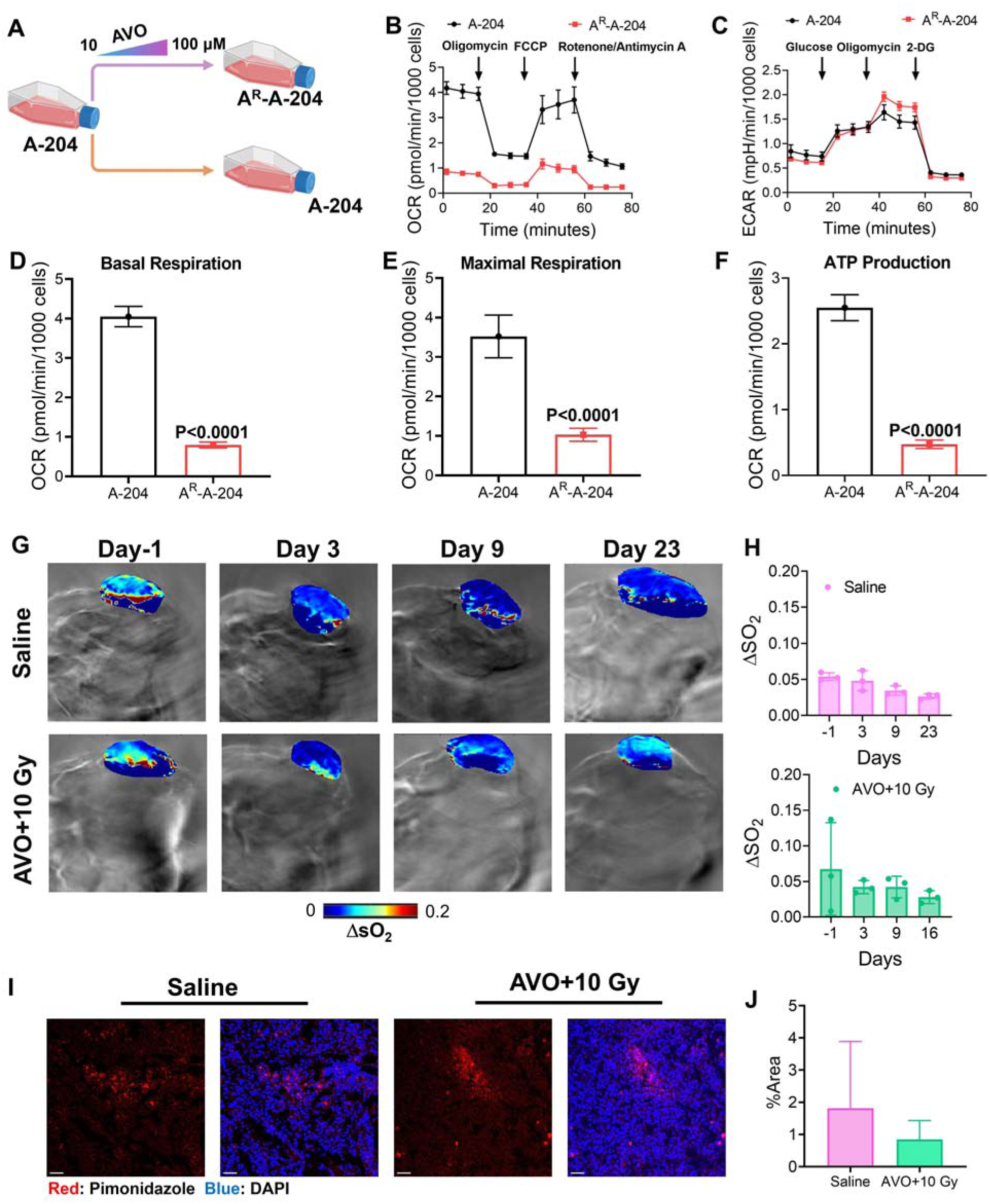
AVO resistant A^R^-A-204 tumors depict altered metabolism and intratumoral hypoxia. **(A)** Scheme depicting the generation of AVO-resistant (A^R^-A-204) cells. **(B)** OCR and **(C)** ECAR traces of wildtype A-204 and A^R^-A-204 cells. **(D, E, F)** Statistically significant reductions in basal respiration, maximal respiration, and ATP production as observed in A^R^-A-204 cells when compared to widtypel A-204 cells. **(G)** Parametric maps and **(H)** quantitative analysis of tumor ΔsO_2_ for saline and AVO+10 Gy-treated A^R^-A-204 xenografts on Day-1, Day 3, Day 9 and Day 23. (n=3). **(I)** Confocal imaging and **(J)** quantitative analysis of pimonidazole staining of excised saline and AVO+10 Gy-treated tumors at endpoint. Scale bar=50 µm.

Next, A^R^-A-204 xenografts were treated with saline or AVO followed by 10 Gy X-Ray and tumor oxygenation was monitored longitudinally using OE-MSOT. As seen in **Figs. 8G** and **H**, saline-treated A^R^-A-204 tumor exhibited higher oxygenation levels at all time points (ΔsO_2_ = 0.048 ± 0.01 at Day 3; ΔsO_2_ = 0.035 ± 0.01 at Day 9 and ΔsO_2_ = 0.027 ± 0.003 at Day 23) when compared to saline-treated A-204 tumors (**Fig. 3C**; ΔsO_2_ = 0.030 ± 0.011 at day 3, ΔsO_2_ = 0.030 ± 0.007 at day 9 and ΔsO_2_ = 0.016 ± 0.01 at Day 23). This finding suggests that constitutive downregulation of OXPHOS in A^R^-A-204 results in less severe hypoxic microenvironment in MRT. Lower pimonidazole immunofluorescence signals (**Fig. 8I** and **Fig. 8J**) validated our *in vivo* OE-MSOT observations. These encouraging data indicate the potential of ΔsO_2_ to serve as an imaging biomarker of treatment resistance to metabolic modulatory therapies.

Further, in the AVO+10 Gy-treated A^R^-A-204 xenografts **(Fig. 8G** and **Fig. 8H)**, OE-MSOT at midpoint revealed no difference from saline treated groups and a significant decrease in ΔsO_2_ (Day 3; 0.042 ± 0.01), contrasting with the A-204 xenografts (**Fig. 3E**; Day 3; 0.142 ± 0.02) which confirms that A^R^-A-204 xenografts no longer respond to AVO treatment and hypoxia alleviation seen in wildtype A-204 tumors was solely reliant on AVO-mediated OXPHOS inhibition. Pimonidazole exhibited an obvious higher signal in the AVO+10 Gy group (**Fig. 8I**) compared to the wildtype A-204 combination treatment group, further supporting our OE-MSOT observations.

### Atovaquone-resistant A-204 xenografts do not respond to AVO-sensitized radiotherapy

Since AVO treatment did not alleviate hypoxia in the A^R^-A-204 xenografts, we further investigated and validated whether the treatment response of AVO+10 Gy would also be diminished in these tumors. As shown in **Fig. 9A**, there was no significant fluctuation in body weight, except for one mouse in the AVO+10 Gy-treated group, which experienced around 25% weight reduction relative to baseline immediately after receiving 10 Gy irradiation. Therefore, this mouse was excluded in the subsequent study. The survival rates and tumor volume changes (**Fig. 9B** and **Fig. 9C**) showed that the average time required for tumors to reach 1000 mm^3^ was 40 days for the saline-treated group. The tumor growth was markedly slower compared to the 29 days (to reach endpoint) observed in saline treated naive A-204 tumors, suggesting that OXPHOS downregulation could make MRT less aggressive. Mice treated with AVO + 10 Gy combination treatment, took an average of 54 days to reach 1000 mm^3^. Although the tumor volume showed a slight reduction after the treatment on Day 9, the tumor continued to grow in the following weeks, indicating A^R^-A-204 tumors did not benefit from the therapy (**Fig. 9C** and **Fig. 9D**). Our data suggests that as expected, AVO no longer exerted a radiosensitizing effect in drug-resistant xenografts. Finally, as shown in **Fig. 9E, H&E** staining showed no significant toxicity observed in the normal organs including heart, liver, kidney and muscle and no detectable morphological changes or apparent necrosis was observed in the tumors excised from either saline or combination therapy cohorts.

**Fig. 9.**
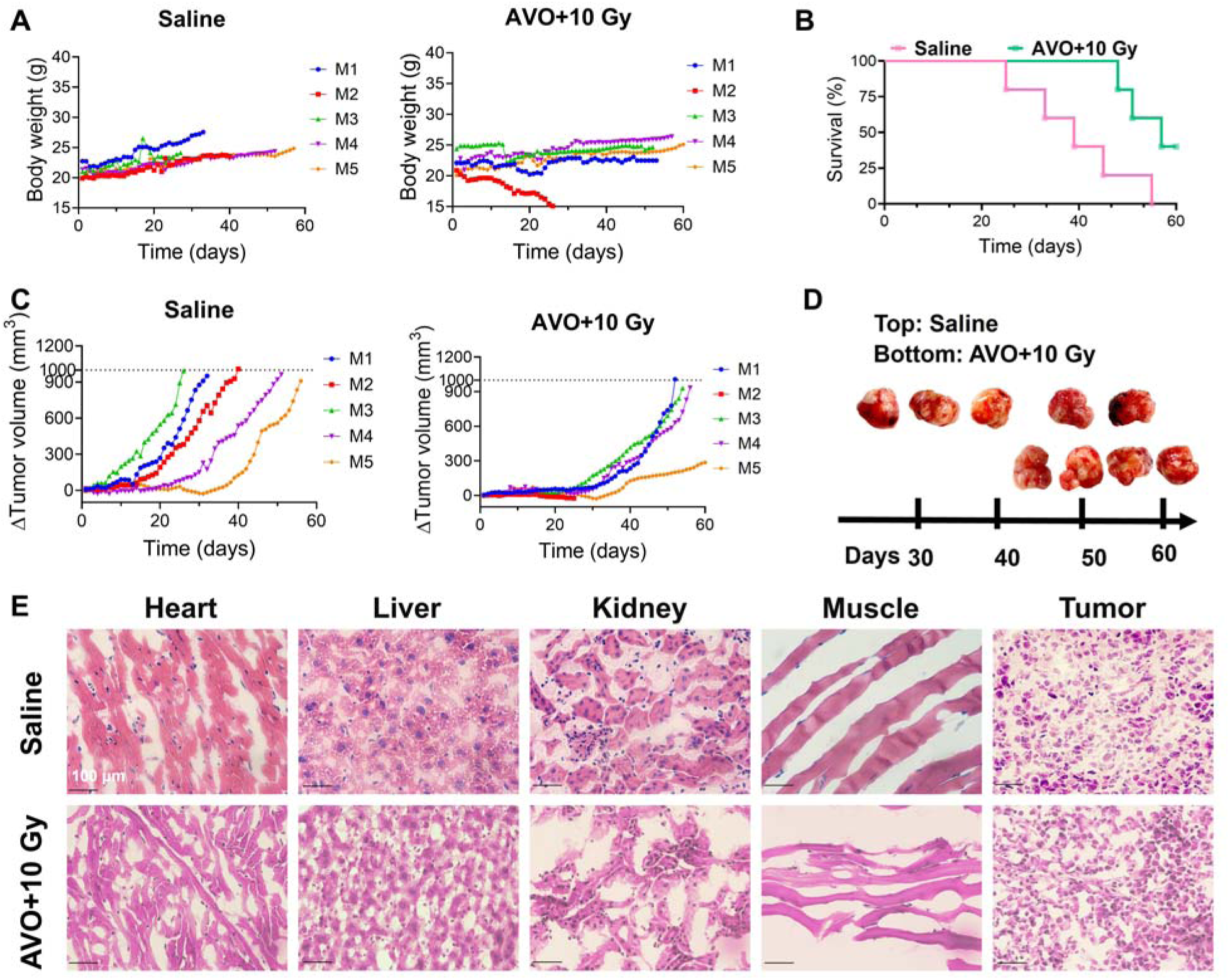
A^R^-A-204 xenografts do not respond to combination therapy. **(A)** Body weight curves of saline and AVO+10 Gy-treated groups. **(B)** Kaplan-Meier survival curves and (**C**) individual tumor growth curves depicting change in tumor volume from baseline, for both groups. (**D**) Photographs of excised tumors taken at endpoint. (n = 5) **(E)** H&E staining of heart, liver, kidney, muscle and tumor tissues of each treamtment groups. Scale bar=100 µm.

### Mathematical modeling reveals a synergistic relationship between AVO and Radiotherapy

A mathematical framework was built to elucidate the dynamic interplay between tumor growth, radiotherapy, and hypoxia alleviation with AVO in MRT. The model was calibrated by fitting it to individual mouse tumor growth kinetics under different treatment conditions. The parameter estimates derived via nonlinear least squares fitting are presented in **Table S1**. As shown in **Figures. S12A–C**, tumor growth kinetics under AVO monotherapy closely resemble those of untreated controls. The estimated tumor growth rate parameter (*γ*) remained effectively unchanged between control and AVO-only conditions (**Table S1**), reinforcing the conclusion that AVO alone did not inhibit tumor progression. When combined with single 10 Gy dose, AVO produced a more pronounced reduction in tumor growth than radiation alone, confirming its role as a radiosensitizer. A strong Pearson correlation between modeled and observed tumor volumes (**Fig. S12, lower panel**) supported the accuracy and robustness of the model fits. These results collectively validate our model’s ability to capture key tumor growth dynamics under varying therapeutic strategies, and they establish a solid foundation for subsequent analyses aimed at optimizing treatment regimens.

Tumor static curve (TSE) analysis which quantifies the relationship between AVO dose, and the corresponding RT dose required to maintain tumor stasis in mice, as obtained from **Equation 1** is presented in **Fig. 10A**. Mathematically, stasis occurs when tumor growth is exactly balanced by treatment-induced cell death, resulting in no net change in tumor volume over time 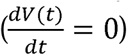 (*42–44*). The TSE curve exhibits a concave-up shape, indicating that the reduction in radiation dose requirements accelerates with increasing AVO doses, particularly in the lower dose range, reflecting a synergistic interaction between AVO and radiation, where AVO enhances radiosensitivity by alleviating hypoxia. At higher AVO doses, the curve begins to flatten, reflecting saturation of this synergistic effect. Above the TSE curve, the shaded green region represents combinations of AVO and radiation doses that not only maintain stasis but also drive tumors into regression. Conversely, dose combinations below the curve are insufficient to counteract tumor growth, resulting in continued expansion of tumor volume. Thus, while the curve itself delineates the boundary between stable and growing tumors, the shaded area highlights the potential for more aggressive tumor shrinkage when both AVO and radiation doses are combined effectively. These findings reinforce the value of AVO as a radiosensitizer and provide actionable insights into dose levels that can shift the treatment goal from stabilization to active tumor shrinkage.

**Fig. 10.**
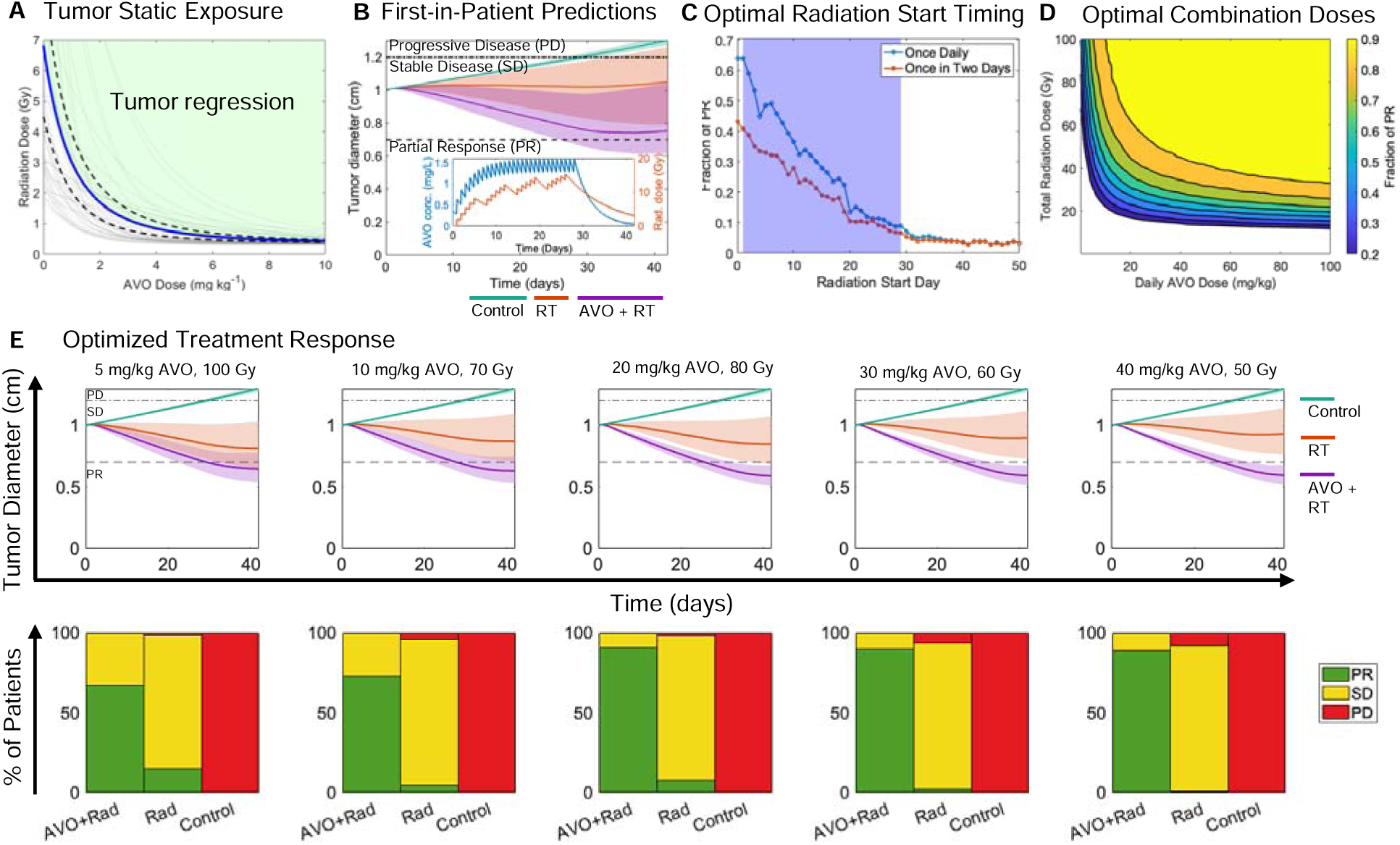
Mathematical modeling predicts efficacy of optimized combination treatment in virtual patient cohort. **(A)** Tumor Static Exposure (TSE) curve (blue) illustrating how increasing AVO dose reduces the radiation dose required for tumor stasis in mice; 95% confidence interval around the TSE curve is shown with dashed black lines; the green area above the curve indicates doses that lead to tumor regression. **(B)** First-in-patient simulations using a regimen of 10 mg/kg/day AVO for 4 weeks and fractionated radiation (2 Gy per fraction, 5 days/week for 4 weeks) show mean tumor trajectories (solid lines) with shaded 95% confidence intervals for control, radiation-only, and AVO+RT treatments in virtual patients. Horizontal dashed and dotted-dashed lines mark thresholds delineating Partial Response (PR), Stable Disease (SD), and Progressive Disease (PD). The inset in **(B)**, displays AVO concentration (left y-axis) and radiation effects (right y-axis) over the extended, fractionated schedule. **(C)** Radiation start time optimization relative to the 4-week AVO schedule (blue band marks AVO treatment period) illustrates how adjusting radiation initiation affects the fraction of patients achieving PR. Simulations included daily AVO administration and an alternate schedule of once every two days to evaluate the effect of dosing frequency on timing optimization. **(D)** Combination dose optimization contour plot shows the fraction of PR across varying AVO and total radiation doses, identifying clinically relevant dose pairs that improve efficacy while reducing radiation exposure. **(E)** Optimized treatment responses at selected AVO/radiation dose combinations: top panels show mean tumor trajectories (solid lines) with 95% confidence intervals for control, radiation-only, and AVO+RT, while bottom panels present bar plots of treatment outcome distributions (PR, SD, PD) within the virtual patient population.

### Translational modeling predicts efficacy of the combination treatment in first-in-human settings

To assess the clinical efficacy of our combination treatment, we applied the mathematical model to a virtual patient cohort to predict tumor response in a first-in-patient setting. (**Fig. 10B**) Parameter values were extrapolated from preclinical mouse data to humans using allometric scaling, while population variability was captured through a virtual patient cohort of 500 individuals. The simulations evaluated tumor dynamics under three treatment conditions: control, radiation-only (2 Gy per fraction, 5 days/week for 4 weeks), and AVO + radiation (10 mg/kg/day AVO for 4 weeks combined with the same radiation schedule). Mean tumor trajectories (solid lines) and 95% confidence intervals (shaded regions) are shown to represent population-level variability in response. Thresholds for Progressive Disease (PD), Stable Disease (SD), and Partial Response (PR), as defined by the clinical gold standard RECIST 1.1 criteria, were delineated by horizontal dashed and dotted-dashed lines. These thresholds directly link the simulated tumor diameter changes to clinical oncology endpoints, enabling the translation of model predictions into meaningful clinical outcomes. The inset in **Fig. 10B** provides additional context, displaying the pharmacokinetic profile of AVO (left y-axis) and the cumulative biological effects of fractionated radiation (right y-axis) over the 4-week treatment schedule.

The results reveal that untreated tumors exhibit continuous growth, while radiation monotherapy slows tumor progression. In contrast, the combination therapy leads to significant tumor shrinkage, demonstrating the radiosensitizing effect of AVO. Of note, the doses used in this numerical experiment were chosen based on existing clinical regimens for other indications and tumor types, rather than being optimized specifically for the context modeled here. This raises the possibility that better outcomes could be achieved through careful optimization of AVO and radiation doses. Accordingly, we undertook a series of optimization experiments.

To assess the impact of radiation start timing relative to AVO and the administration frequency of AVO on treatment outcomes, simulations were performed for radiation initiation days ranging from day 0 (simultaneous with AVO start) to Day 30 (radiation starting 30 days after AVO initiation). AVO was administered at a dose of 10 mg/kg either daily or once every two days for four weeks, while radiation was delivered at 2 Gy per fraction, five days per week, for four weeks.. **Fig. 10C** demonstrates that daily AVO administration leads to significantly higher PR rates across all start times compared to the alternate once-every-two-days schedule. The highest PR rates are observed when radiation is initiated concurrently with or shortly after the start of AVO treatment, emphasizing the importance of synchronizing radiation delivery with the radiosensitization effects of AVO. Delaying radiation initiation within the 4-week AVO treatment window results in a steep decline in PR rates. Beyond the 4-week AVO treatment period, PR rates plateau at persistently low levels, reflecting the diminished radiosensitization effect once AVO administration ends. These findings highlight the necessity of maintaining consistent AVO plasma concentrations and synchronizing radiation timing during the active treatment phase to maximize the synergistic effects of the combination therapy.

Further, **Fig. 10D** explores the synergistic effects of AVO and radiation dose combinations by simulating 2,500 dose pairs (50 values each for AVO and radiation doses) and calculating the fraction of patients achieving PR for each combination. AVO doses ranged from 0.1–100 mg/kg/day, while total radiation doses ranged from 0.1–100 Gy (administered as fractionated doses over 4 weeks). The resulting contour plot demonstrates that higher AVO doses significantly reduce the radiation required to achieve PR in a substantial proportion of the virtual patient population, underscoring the synergistic interplay between the two modalities. Notably, in corroboration with the TSE analysis, diminishing returns at very high AVO doses were observed, as reflected in the large horizontal spread of the yellow region, where further increases in AVO provide minimal incremental benefit in reducing radiation requirements.

Finally, we evaluated the impact of optimized treatment regimens on tumor outcomes in the virtual patient cohort. The five regimens analyzed were selected from the contour plot (**Fig. 10E**) to represent diverse combinations of AVO and radiation doses, spanning low AVO with high radiation (e.g., 5 mg/kg AVO + 100 Gy) to high AVO with low radiation (e.g., 40 mg/kg AVO + 50 Gy). The top row of **Fig. 10E** presents mean tumor trajectories (solid lines) with 95% confidence intervals (shaded regions) under each regimen, alongside control and radiation-only treatments. The bottom row displays the corresponding outcome distributions as bar plots, categorizing the percentage of patients achieving PR, SD, or PD. A clear enhancement in treatment efficacy as AVO doses increase and RT doses decrease across the selected regimens can be seen. Lower-dose regimens (e.g., 5 mg/kg AVO + 100 Gy) result in modest tumor shrinkage, with a higher proportion of patients classified as SD. As AVO doses increase (e.g., 40 mg/kg AVO + 50 Gy), tumor shrinkage becomes more pronounced, reflected by a larger fraction of patients achieving PR and a marked reduction in SD cases. This highlights the potential of AVO to reduce the reliance on high radiation doses while maintaining or even improving treatment outcomes. Notably, the combination regimens show a clear advantage over RT monotherapy, which achieves only modest tumor control. Incorporation of AVO not only improves PR rates but also reduces the proportion of patients with SD or PD, shifting the outcome distribution more favorably toward tumor shrinkage across the virtual patient cohort.

## Discussion

Over the past two decades, multiple therapeutic approaches, such as surgery, chemotherapy and radiation, have been employed to enhance treatment outcomes and survival rates in pediatric MRT and ATRT patients. RT remains the mainstay of treatment for both cranial and extracranial rhabdoid tumors. However, despite major efforts, significant advances remain elusive, especially in patients less than 3 years old due to long-term toxicities from the high doses of RT and chemotherapy required to achieve meaningful responses (*4, 45*). Thus, there has been an increasing interest in exploring novel therapeutic strategies to maximize RT efficiency at lower doses to minimize immediate and long-term radiotoxicity in pediatric MRT management. However, new drug discovery is scientifically and financially challenging for rare tumors like MRT and ATRT. Thus, drug repurposing is an attractive strategy. AVO is an FDA approved antimalarial drug and has been reported for the treatment of various adult malignancies including breast cancer (*23*), renal cell carcinoma (*46*), non-small cell lung cancer (*22*), aggressive thyroid cancer (*47*) and leukemia (*48*), both as monotherapy and in combination with chemoradiation. However, to the best of our knowledge, this is the first work utilizing AVO to enhance radiosensitivity in pediatric cancer treatment and management.

Our findings in this study unequivocally demonstrate the benefit of combining AVO with RT to achieve complete, durable therapeutic responses in MRT xenografts, at low, clinically viable doses. Our results revealed that short term (7 day) AVO treatment at pharmacologically viable doses could effectively improve the oxygenation levels in A-204 tumors (**Fig. 3**), and thereby result in long-term inhibition of tumor growth when combined with a single 10 Gy X ray dose. (**Fig. 5**) This phenomenon was confirmed by our histopathological data. Additionally, compared to the saline-treated group, the lower Ki-67 signal (**Fig. 4**) and higher γ-H2AX signal (**Fig. 6**) further validated that the combination therapy enhanced DNA damage and cell apoptosis, ultimately inhibiting the tumor growth. Most strikingly, for the AVO+RT treatment groups, all the tumors were completely abolished within 10 days post-treatment, and did not recur for the remainder of the study. Our approach demonstrated exceptional treatment outcomes when compared to other MRT treatment methods, as most existing strategies can only inhibit tumor growth without fully eliminating them (*49, 50*). showing that a negligible pimonidazole signal was observed in the AVO-treated groups (**Fig. 4**).

There are several advantages to our treatment strategy: (1) Although multiple treatment strategies including immunotherapy and targeted therapy, have been explored in recent years to improve the survival rates of MRT patients, conducting comprehensive studies of these novel treatment strategies remains challenging due to the rarity of pediatric MRT and limited patient samples. Thus, the traditional combination of chemotherapy and radiation therapy for treating MRT patients would be the quickest approach to investigate further and implement in clinical practice; (2) Before a new drug can be administered to clinical MRT patients, several critical steps must be completed, including preclinical studies, an Investigational New Drug (IND) application, clinical trials (Phases I-III), a New Drug Application (NDA), regulatory approval, and post-market surveillance (*51*). Given that most patients with rhabdoid tumors are under 3 years old, using an orally bioavailable, FDA approved, off-label drug, AVO to radiosensitize MRT in the clinic can be easily and quickly envisaged, as opposed to other radiosensitizers currently under development that would need to undergo a long-term approval. Additionally, our data demonstrates that AVO exhibits effects at significantly lower concentrations (EC_50_ = 28.36 µM, **Fig. 2A**) than that of Metformin (EC_50_ = 13.15 Mm, **Fig. S1**) and another complex I inhibitor, phenformin (*52*). Therefore, although safe for use, these biguanides have resulted in suboptimal efficacy in clinical trials. Another preclinical inhibitor of mitochondrial respiratory complex I, IACS-010759, has been associated with multiple side effects such as neurological symptoms (*53*), hematologic toxicity, metabolic disturbances, weight loss and nausea and vomiting (*54*), owing to extremely narrow therapeutic indices. Both phenformin and IACS-010759 were withdrawn from clinical trials due to a risk of fatal lactic acidosis (*52,53*) resulting from a shift towards glycolysis that induces increased lactic acid production and a heightened risk of fatal lactic acidosis. Our preliminary metabolic analysis does not indicate an obvious glycolytic shift in A-204 cells even after prolonged AVO exposure (**Fig. 8** and **Fig. S2**) which bodes well for its translation to pediatric patients. Moreover, our data monitoring mouse body weight (**Fig. 5B**) and histopathology of off-target organs (**Fig. 6D**) demonstrates that AVO does not cause noticeable systemic toxicity. Additionally, as an off-label drug, cost-effectiveness of AVO can double the impact to patients suffering from this disease, especially considering the small patient cohorts. Therefore, considering all the benefits, AVO stands out as the optimal mitochondrial inhibitor in pediatric MRT management; (3) Local radiation therapy was employed in this work. This targeted and precise treatment allows for highly accurate radiation delivery, reducing side effects and enhancing treatment outcomes by focusing directly on the tumor while minimizing damage to surrounding health tissues. Our data establishes that AVO can integrate both with single dose and fractionated RT regimens with ease and without changes to its effectiveness. Most excitingly, AVO reduces the effective local radiation dose required to achieve maximum and durable therapeutic effect, which makes RT safer for pediatric patients both in the long and short term.

Another unique feature of our work is the adoption of noninvasive MSOT imaging to map the spatiotemporal changes in pediatric rhabdoid tumors, in response to AVO. Our imaging results suggest that the real-time changes in oxygenation could be visualized and monitored before, during, and after the treatment with high fidelity and reproducibility. MSOT is a safe, widely available, and ionizing radiation-free imaging modality, which can inform tumor oxygenation status in a label-free manner. By performing %sO_2_ measurement following a switch of 21% oxygen to 100% oxygen, using the oxygen-enhanced (OE-MSOT) procedure, ΔsO_2_ was be measured. Our data indicates the potential of ΔsO_2_ to serve as both a pharmacodynamic biomarker measuring the magnitude and duration of hypoxia resolution by AVO, as well as a biomarker to indicate the onset of resistance to the drug. We show that although potent, hypoxia alleviation by AVO is transient, which necessitates careful timing of RT. As it is free from ionizing radiation, we posit that MSOT can be easily and safely implemented to guide and optimize treatment schedules, as well as evaluate treatment response of AVO-enhanced RT in pediatric MRT patients, enabling real-time, personalized treatment adjustments. These studies fill an unmet gap in the design and implementation of smart, data-driven combination therapies, particularly for children and young adults where poorly designed treatments can lead to devastating acute and chronic toxicities. OE-MSOT has the potential in refining the administration of anticancer therapies, especially by optimizing the timing of sequential treatments and distinguishing between responsive and unresponsive tumors. Furthermore, frequent monitoring enabled by an ionizing radiation-free imaging modality would spare the pediatric patients from ineffective or toxic over-treatment. Although clinical trials with AVO are currently underway in adult cancers (*22*), the results of our study provide a strong foundation for advancing the drug to benefit children with rhabdoid tumors and other radioresistant cancers.

Finally, to extrapolate our preclinical findings to human patients, we have developed innovative mathematical models to predict human responses as well as determine optimal AVO+RT dosing and timing combinations. By bridging preclinical and clinical settings, these first-in-patient simulations underscore the model’s utility for predicting patient-specific outcomes and optimizing treatment strategies. The inclusion of inter-patient variability in the virtual population ensures robustness of the predictions, supporting future clinical trial designs that aim to refine dosing regimens and schedules based on expected clinical response rates. By integrating both tumor response trajectories and categorical outcome distributions, our data (**Fig. 10E**) provides a comprehensive view of how optimized regimens can shift the balance toward more favorable outcomes in a virtual patient cohort. These findings reinforce the importance of tailoring dosing strategies to maximize treatment efficacy while minimizing the potential for radiation-induced toxicity.

Our mathematical model has some limitations that warrant consideration. First, the inability of the model to achieve CR is rooted in its mathematical structure. By design, the model describes tumor dynamics using continuous differential equations, where tumor volume can asymptote toward zero but never reach it exactly. As a result, any residual tumor volume, however small, persists at the end of the simulation and can regrow over time, preventing complete regression. To address this, one could redefine CR based on a practical threshold, such as a 99% reduction in tumor volume. However, for simplicity, we chose to focus on PR, which is a clinically relevant metric aligned with RECIST 1.1 guidelines. The translational relevance of the findings would also benefit from further validation using clinical datasets. The optimization studies explored fixed dose schedules, but adaptive treatment strategies and considerations of healthy tissue toxicity remain areas for future exploration. Finally, we recognize that children and infants are not small adults, and the differences are not merely body weight-dependent, but rooted in physiological and biochemical differences (*55*). Thus, for simplicity, our virtual patient cohort is generated taking an adult human patient into account (70 kg), which may not fully capture the complexities of translating the combination therapy in the pediatric setting. Addressing these limitations in future studies will enhance the model’s applicability and translational value.

## Materials and Methods

### Data Acquisition and Processing of mRNA Sequencing Expression Profiles

We downloaded the transcriptomic data from National Cancer Institute (NCI) Therapeutically Applicable Research to Generate Effective Treatments -Rhabdoid Tumor (TARGET-RT) project (https://portal.gdc.cancer.gov/projects/TARGET-RT) within the Genomic Data Commons (GDC) Data Portal (Data Release 37.0-March 29, 2023) using The Cancer Genome Atlas (TCGA) biolinks R package (https://bioconductor.org/packages/release/bioc/html/TCGAbiolinks.html). Raw counts and sample metadata were loaded in the R environment using the Huntsman Cancer Institute (hciR) package (https://github.com/HuntsmanCancerInstitute/hciR) of the University of Utah. 63 rhabdoid cancer patients, all aged between 0 and 14 years were selected. Log2 normalization was performed for further analysis. Single sample gene set enrichment analysis (ssGSEA) was performed based on Hallmark gene sets. Enrichment scores for all pathways are plotted in a heatmap, categorized by the different stages of rhabdoid tumor patients.

To plot survival curves, we first loaded the R package – survival: survival analysis (ver 3.7-0) and ran the Cox proportional hazards regression modeling on all samples. We then looped through all 230 OXPHOS genes (OXPHOS gene signature; KEGG database), applying the Cox proportional hazards model to obtain the Cox coefficients and p-values. Next, we ranked the OXPHOS genes and identified those with significant p-values (p < 0.05), indicating a difference in survival rates based on high or low gene expression. Finally, we plotted survival curves for these genes by dividing the survival data into high and low expression groups based on the median and included the p-value in the plot.

### Cell lines and Compounds

A-204 cell line was purchased from the American Type Culture Collection (ATCC). Cells were cultured at 37°C with 5% CO_2_ in McCoy’s 5A medium (ATCC) supplemented with 10% fetal bovine serum (FBS, GenClone) and 1% penicillin-streptomycin (GenClone). Atovaquone was obtained from Thermo Scientific. Bovine serum albumin (BSA) was obtained from Sigma. Matrigel was obtained from Huntsman Cancer Institute Preclinical Research Resource. Pimonidazole hydrochloride and Red 549 Mab were obtained from Hypoxyprobe^TM^. Purified Rat Anti-mouse CD31 (Catalog No: 550274) was purchased from BD Biosciences. Alexa Fluor 488 AffiniPure Donkey Anti-Rat lgG(H+L) (Lot 163136) and Rhodamine (TRITC) AffiniPure Donkey Anti-Rat lgG(H+L) (Lot 157518) were obtained from Jackson ImmunoResearch Inc. Histone H2AX antibody (NB100-384) was purchased from Novus Biologicals and secondary antibody Donkey anti-Rabbit lgG (H+L) Dylight594 (SA5-10040) from Invitrogen. Purified anti-mouse/human Ki-67 antibody (151202) was purchased from Biolegend. The 2,5-diphenyltetrazolium bromide (MTT) assay (M6494) was obtained from ThermoFisher Scientific.

To model acquired resistance, atovaquone-resistant A-204 cell line (A^R^-A-204) was established in house by gradient AVO treatment following slight modifications of previously published protocols (*56*).

### Cell Viability Assay

To determine the cytotoxicity of AVO, MTT assay was carried out in A-204 and A^R^-A-204 cells. The cells were seeded at a density of 2000 cells/well in a 96-well plate and cultured in McCoy’s 5A culture medium at 37°C with 5% CO_2_. After 24 hours incubation, cells in each well were treated with various concentrations of AVO (5, 10, 20, 50, 100, 200 and 500 µM) in 100 µL culture medium, with each concentration tested in 6 replicates. Following 72 hours of AVO treatment, MTT solution (10% 12 mM MTT stock solution) was added to each well and incubated for 4 hours. The medium was then removed, and 50 μL of DMSO was added to each well. The cells were incubated for an additional 10 minutes at 37°C. The absorbance at 540 nm was determined using a microplate reader (Cytation 3, BioTek, USA) and the cell viability was calculated using the following equation:

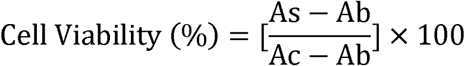

As = Absorbance of Test well;

Ab = Absorbance of Blank (medium only);

Ac = Absorbance of Control (cells only)

### Seahorse Experiment

Real-time oxygen consumption rates (OCR) and extracellular acidification rates (ECAR) for A-204 and A^R^-A-204 cells treated with AVO were determined using the XFPro Analyzer (Seahorse Biosciences). A-204 and A^R^-A-204 cells were seeded in the Seahorse XF Pro cell culture microplate at a density of 20,000 cells/well and cultured at 37°C with 5% CO_2_ in McCoy’s 5A medium for 24 hours to allow attachment. Then, both A-204 and A^R^-A-204 cells were treated with 25 µM AVO for 24 hours. Control groups treated with DMSO alone were processed in parallel. The measurements of the OCR were taken in XF assay medium containing 10 mM glucose. 1 mM pyruvate and 2 mM L-glutamine and 1 µM oligomycin, 0.5 µM phenyl-hydrazone (FCCP) and 0.5 µM rotenone/antimycin A were added sequentially for OCR measurement. ECAR measurements were taken in XF assay medium containing 1 mM pyruvate and 2 mM L-glutamine. 10 mM glucose, 2 μM oligomycin and 100 mM 2-DG were added for ECAR into the injection ports in the XF Pro sensor cartridge. 25 μM AVO was added into each well (excluding the DMSO wells) for the duration of the seahorse experiment. Measurements were normalized cell numbers, counted by adding 2 μM Hoechst 33342.

### Animal Models and Treatments

Mice were purchased from Charles River Laboratories. All animal experiments were conducted in accordance with the approved protocol by the Institutional Animal Care and Use Committee (IACUC) at the University of Utah. Wildtype A-204 (1 × 10^6^) cells or AVO-resistant A^R^-A-204 (1 × 10^6^) cells (in PBS: Matrigel=1:1 solution) were inoculated subcutaneously in 26-44 days old athymic nude female mice. When the tumors reached around 50 mm^3^, mice were randomized into treatment groups (n=5) that were given saline (control), AVO, RT (10 Gy) or the combination treatment of AVO followed by RT. AVO was administered as an oral suspension of 50 mg kg^-1^ dose per day via oral gavage, following clinical dosing schedule (*57*). Mice were given this dose for 7 days because AVO typically requires one or two weeks to achieve a consistent plasma concentration in human patients (*58, 59*). Mice in RT or combination treatment cohorts received a single dose of 10 Gy X-ray irradiation focused on the tumor (Rad Source Technologies, USA) after 7 days of AVO treatment. A separate cohort of mice was also treated with 30 Gy dose either alone (in fractions of 6 Gy over 5 days) or in combination with AVO.

Mice were monitored for welfare throughout the study, and body weights were collected every day. Tumor size was measured with calipers and volume was calculated with the formula: [V=(W^2^×L)/2]. Mice were euthanized (study endpoint) once the tumor reached 1000 mm^3^, or if the body weight dropped 20% relative to baseline or if the mice were under poor bodily condition. After sacrifice, tumors and other organs were excised 1 hour after intraperitoneal (i.p.) administration of Pimonidazole hydrochloride (60 mg kg^-1^). A^R^-A-204 tumor-bearing mice were subjected to the same procedures.

### In vivo MSOT

All *in vivo* MSOT experiments were performed using iThera inVision 256-TF MSOT System (iThera Medical GmBH). MSOT scans were performed 1 day prior treatment (referred to as Day-1 when the tumors reached to ∼ 50 mm^3^) for all groups to establish baseline. Subsequently, for the saline and AVO-treated groups, longitudinal MSOT was conducted on Day 3, Day 9 and Day 23 post-treatment. For the 10 Gy and AVO+10 Gy-treated groups, MSOT scans were conducted on Day 3, Day 9, Day 16 and Day 30.

Briefly, A-204 or A^R^-A-204 tumor-bearing mice were anesthetized using isoflurane and tail veins were catheterized. A thin layer of ultrasound gel (Aquasonic Clear, Parker Labs) was applied to facilitate optical and acoustic coupling with the polyethylene membrane. Subsequently, the mice were positioned in a lateral, side-lying orientation in a bag for imaging purposes. The chamber temperature was adjusted to 36 °C and the mice were allowed to equilibrate to the temperature for 10 minutes before imaging. This temperature allowed the mice to maintain 37 °C body temperature for the duration of the scan. Images were acquired at the largest tumor cross section of the mice with a step size of 1 mm. OE-DCE MSOT imaging was performed in the same scan session, following our previously published protocols (*30*). For OE-MSOT scan, images were first acquired at 700 nm, 730 nm, 760 nm, 800 nm, 850 nm, and 875 nm wavelengths with 21% O_2_ (air) breathing gas for 2 minutes. 10 frames and 20 repetitions were recorded at every imaging wavelength. The breathing gas was switched to 100% O_2_. To allow the mice to reach equilibrium, the mice remained in the chamber for 2 minutes without imaging. A second set of images were collected with 100% O_2_ breathing gas for 2 minutes using the same parameters.

For DCE-MSOT scans, images were acquired at 700 nm, 730 nm, 760 nm, 800 nm, 850 nm, and 875 nm wavelengths under 100% O_2_ breathing gas. 10 frames and 141 repetitions were recorded at every imaging wavelength. After 1 minute of acquiring pre-injection baseline images, indocyanine green (ICG, Sigma, 300 nmol, 100 µL) was intravenously injected and mice were scanned continuously for additional 15 minutes. ViewMSOT (Version_4.0), was utilized for image processing. Images were reconstructed using the Model Linear algorithm and fluence corrected (only for OE-MSOT scans) according to vendor presets. Spectral unmixing extracted the relative contribution of oxyhemoglobin (HbO_2_) and deoxyhemoglobin (Hb), which were then utilized to produce blood oxygen saturation measurements (%sO_2_ ^air^ and %sO_2_ ^O2^) with each breathing gas. The difference between the average %sO_2_^air^ and %sO_2_^O2^ during their respective 2 minutes acquisitions was then used to determine ΔsO_2_ within the tumor. For the DCE-MSOT scan, the pharmacokinetic rates of uptake (NK^trans^) and wash-out (K_ep_) of ICG were computed using our fluence-independent image analysis protocol (*30, 60*). All parametric analyses for OE-DCE-MSOT scans were performed in MATLAB R2021b using our custom code (available upon request).

### Histological Analysis

Tissues prepared for immunofluorescence were rinsed in PBS and preserved in OCT (Fisher HealthCare, USA) at −80°C. Tissues were cryo-sectioned into 5 µm thick sections (ARUP Laboratories, University of Utah) and stored at −80°C for further use. Prior to use, tissue sections were fixed with cold 4% PFA for 10 min and washed twice in PBS, followed by incubation in blocking buffer (5% BSA in PBS supplemented with 0.1 % Tween 20) for 1 hour. For pimonidazole staining, sections were stained with Red 549 dye-MAb (1:100, diluted in blocking buffer) overnight at 4°C and thoroughly washed twice in PBS. For CD31, γ-H2AX and Ki-67 staining, tissues were first incubated overnight at 4°C with primary rat anti-mouse CD31 (1:100, diluted in blocking buffer), primary rabbit γ-H2AX (1:100, diluted in blocking buffer) or primary rat anti-mouse/human Ki-67 (1:100, diluted in blocking buffer), respectively. After washing twice with PBS, the sections were incubated for 1 hour at room temperature with secondary antibodies: Alexa Fluor 488 Affinipure Donkey Anti-Rat lgG (H+L) (1:200, diluted in blocking buffer), Dylight594 Donkey Anti-Rabbit lgG (H+L) (1:200, diluted in blocking buffer), or TRITC Affinipure Donkey Anti-Rat lgG (H+L) (1:200, diluted in blocking buffer) for CD31, γ-H2AX and Ki-67, respectively. Sections were washed twice in PBS, stained with 1 µg/mL DAPI for 10 minutes, followed by mounting buffer. All slides were stored at 4°C in a light-protected environment. Fluorescence images were acquired with the confocal laser scanning microscope (CLSM, Leica, SP8). At least 5 separate regions were imaged per tissue and 3 mice per group were analyzed for histological experiments. Whole tissue images were obtained on an Axioscan 7 slide scanner. For histological evaluation of tissues, 5 µm sections were obtained from the OCT cassettes and subsequently utilized for hematoxylin and eosin (H&E) staining. Slides were scanned at 40 × resolution using the Olympus biological microscope (Olympus BX41) and the images were processed using Amscope software.

### Mathematical Model Development

We developed a mathematical framework to elucidate the dynamic interplay between tumor growth, radiotherapy, and pharmacological intervention with AVO in MRT. Expanding upon our prior mechanistic model (*61–63*), the current model integrates tumor growth dynamics, drug pharmacokinetics, and the modulation of oxygen saturation (%sO_2_) within hypoxic tumor microenvironments, thereby capturing the biological processes that influence treatment efficacy.

### Tumor Static Exposure Curve

The Tumor Static Exposure (TSE) curve relates the radiation dose (Dose_rad_) to the AVO dose (Dose_AVO_) required to maintain tumor stasis, defined by *dV*(*t*)/*dt* = 0. (*42–44*) At *dV*(*t*)/*dt* = 0, tumor growth is exactly countered by treatment effects, resulting in a stable tumor size. By examining this curve, we can quantify how increases in AVO dose reduce the radiation dose needed to achieve tumor stasis, thereby providing insights into the synergistic interaction between AVO-mediated oxygenation enhancement and radiation therapy.

Assuming a single dose of AVO and a steady-state approximation for its pharmacokinetics, the relationship between Dose_rad_ and Dose_AVO_ is derived as:

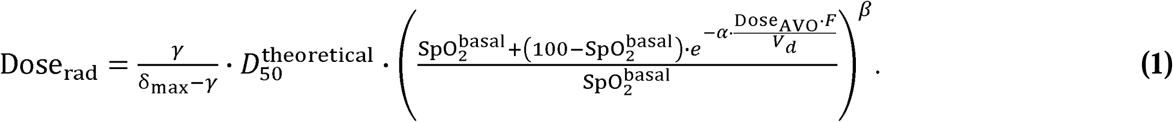

Full derivation details are provided in the Supplementary Information.

To generate the TSE curve, we solved **Equation 1** for 500 values of Dose_AVO_ ranging from 0 to 10 mg/kg, calculating the corresponding Dose_rad_ values for each point. We then repeated this process 50 times, each time randomly sampling parameters from the mouse-based parameter distributions obtained during model calibration. Parameters were sampled using Latin Hypercube Sampling (LHS) to capture inter-animal variability. The mean value and 95% confidence intervals of all simulations were obtained and plotted to illustrate the relationship between Dose_AVO_ and Dose_rad_ .This approach incorporates inter-individual variability and provides a rigorous means to evaluate how varying AVO dosing can reduce the required radiation dose, offering quantitative insights into potential synergistic treatment strategies.

### Interspecies Scaling

To evaluate the translational potential of our framework for predicting AVO pharmacokinetics and tumor response dynamics, we extrapolated the mouse model to humans. This was achieved by substituting mouse-specific parameter values with human population averages from the literature (*64*) or by applying allometric scaling to parameters derived from the mouse model. The scaling was performed as:

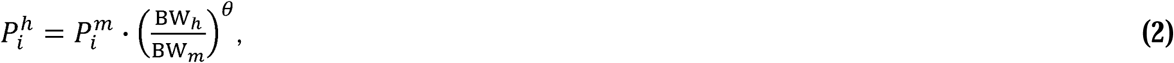

where, 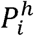 and 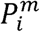 is the value of parameter & for humans and mice, respectively; BW*_h_* and BW*_m_* is the body weight of humans (70 kg) and mice (0.02 kg), respectively. The allometric scaling exponent (*θ*) varies depending on the parameter type. For rate constants such as the tumor growth rate (γ) and maximum radiation-induced cell death rate (δ_max_) was set to –0.25. For parameters related to the drug, including the sensitivity of oxygenation to AVO (*α*) and clearance (Cl), *θ* was set to 0.75. For the volume of distribution (*V_d_*), *θ* was set to 1. This approach ensures a physiologically consistent extrapolation of pharmacokinetic and dynamic parameters from mice to humans, enabling a preliminary assessment of how AVO dosing and tumor responses might translate to clinical contexts.

### Virtual Patient Population Generation

To explore inter-individual variability in treatment response, we generated a virtual patient population by sampling model parameters from distributions derived for humans. Nine key parameters were included in the sampling: *α*, *β*, *γ*, *δ*_max_, 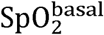, *k*, Cl, *V_d_,* and *F*. The parameter distributions were informed by literature values or estimates obtained from the interspecies scaling process.

LHS was employed to efficiently sample parameter values from these distributions. This method stratifies each parameter’s range into equally probable intervals and ensures comprehensive exploration of parameter space. Using LHS, we generated a total of 500 virtual patients, each characterized by a unique combination of parameter values. This approach captures inter-individual variability in pharmacokinetics, tumor characteristics, and treatment responses, enabling robust evaluation of AVO and radiation therapy under diverse patient scenarios.

### Tumor Response Evaluation Metrics

Tumor response to treatment was evaluated based on percentage changes in tumor diameter relative to baseline, consistent with clinical standards such as the RECIST 1.1 criteria (*65*). In our simulations, the tumor was assumed to be spherical, and the diameter (*d*(*t*)) was defined as the diameter of the sphere at a given time point. The percentage change in tumor diameter (Δ*d*) was calculated as:

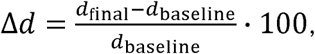

where *d*_baseline_ is the tumor diameter at the start of treatment and *d*_final_ is the diameter at the end of the simulation period (i.e., 30 days after treatment completion). Tumor responses were categorized as follows: Complete Response (CR) was defined as the disappearance of the tumor; Partial Response (PR) was defined as Δ*d* ≤ −30%, representing a reduction of 30% or more in tumor diameter relative to baseline; Stable Disease (SD) was defined as a change of −30% < Δ*d* <+20%; and Progressive Disease (PD) was defined as < Δ*d* ≥ +20% corresponding to an increase of 20% or more in tumor diameter relative to baseline.

This metric provides a clinically relevant evaluation of treatment efficacy by focusing on changes in tumor dimensions rather than volumes. By aligning with clinical standards for tumor response, this method enhances the translational potential of the results.

### Optimization of Treatment Protocols

To identify optimal treatment strategies, we conducted two sets of optimization experiments: (1) determining the best timing for initiating radiation therapy relative to AVO administration and (2) identifying clinically optimal dose combinations of AVO and radiation that minimize total radiation exposure while achieving effective tumor control. Both sets of experiments used the virtual patient population of 500 individuals, and tumor response was evaluated based on percentage change in tumor diameter ( ) one month after the end of treatment, consistent with RECIST 1.1 criteria. The fraction of patients achieving a PR ( ≤ −30%) was used as the primary outcome metric. Additionally, to assess the impact of AVO administration frequency on treatment outcomes, simulations included two AVO dosing schedules: daily and once every two days. The details can be found in the Supplementary Information.

### Statistics

Statistical analysis was conducted using GraphPad Prism 8 software (GraphPad Software Inc.). The data were analyzed using Unpaired *t*-tests and One-Way ANOVA (multiple comparisons). Statistical significance between two groups was determined with two-tailed P-values: *P < 0.05, **P < 0.01, ***P < 0.001, and ****P < 0.0001. Survival was estimated by comparison of Kaplan-Meier curves using a log rank test.

## Acknowledgements and Funding

This work was supported, in part, by the University of Utah College of Pharmacy (S.G.), the University of Utah Immunology, Inflammation and Infectious Diseases (3i) Initiative (S.G. and S.S.), the 5 For the Fight Fellowship (S.G.), and the Elsa U Pardee Foundation (S.G)., the University of Utah school of Medicine (S.S.) and the Office of the Vice President for Research Seed Grant (S.S.) and Research Instrumentation Fund (S.G.) and NIH grant R01EB035545 (P.D.). We acknowledge direct financial support from the Huntsman Cancer Center supported by the National Cancer Institute of the National Institutes of Health under Award Number P30CA042014.

## Author contributions

S.G. and S.S. conceived and designed the experiments, W.X. performed the experiments and data analysis. P.D. conceived and designed mathematical modeling experiments. C.S. and J.C. performed mathematical modeling. M.G. and N.S. analyzed the MSOT imaging data. M.G. designed the TOC. J.H. and E.N. performed *ex vivo* imaging experiments and data analysis. D.G. and R.J. provided assistance with X-ray irradiation. M.D.P. and S.G. created the OE-DCE MSOT MATLAB code. W.X., S.G. and P.D. wrote the manuscript. S.G. and S.S. edited the manuscript.

## Data and materials availability

All data needed to evaluate the conclusions in the paper are present in the paper and/or the Supplementary Materials. Additional data related to this paper may be requested from the authors.

