## Supplemental Data for "Image-Guided Targeting of Mitochondrial Metabolism Sensitizes Pediatric Malignant Rhabdoid Tumors to Low Dose Radiotherapy"

Wenxi Xia *et al.*

**This file includes:** Figs. S1 to S12 and Table S1

**Methods**

**Mathematical Model Development**

The time-dependent evolution of tumor volume ($V\left( t \right)$) is governed by exponential tumor growth and radiation-induced tumor cell death:

$\frac{dV\left( t \right)}{dt}=\gamma\cdot V\left( t \right)-\delta_{\max}\left( \frac{D\left( t \right)}{D_{50}^{\mathrm{effective}}+D\left( t \right)} \right)\cdot V\left( t \right)$,	$V\left( 0 \right)=V_{0}$	**(S1)**

where $\gamma$ is the tumor growth rate, $\delta_{\max}$ is the maximum radiation-induced cell death rate, $D\left( t \right)$ is the radiation dose at time $t$, $D_{50}^{\mathrm{effective}}$ is the oxygen-modulated radiation dose achieving half-maximal tumor cell death, and $V_{0}$ is the tumor volume at the start of treatment. The cell death term incorporates a Michaelis-Menten function, $\frac{D\left( t \right)}{D_{50}^{\mathrm{effective}}+D\left( t \right)},$ to reflect saturation effects at high radiation doses.

The radiation dose $D\left( t \right)$ was modeled as an exponentially decaying function:

$D\left( t \right)=\sum_{j=1}^{M} \mathrm{Dose}_{\mathrm{rad}}\cdot e^{-k\left( t-t_{j} \right)}\cdot H\left( t-t_{j} \right)$,	**(S2)**

where $\mathrm{Dose}_{\mathrm{rad}}$ is the administered radiation dose, $k$ is the biological decay rate constant of radiation efficacy, and $t_{j}$ denotes the time of the $j$-th radiation administration, with $j=1,2,\ldots,M$. The Heaviside function $H\left( t-t_{j} \right)$ ensures that each dose contributes only after its administration. In the present mouse study, a single external beam radiation dose was used, simplifying the summation to one term. In future clinical simulations, fractionated dosing can be modeled by using multiple terms in the summation to capture the cumulative effects of repeated administrations.

The exponential decay reflects the biological effects of radiation, rather than physical dissipation. This decay accounts for processes such as the repair of radiation-induced DNA damage and the redistribution of cellular damage effects over time, providing a computationally efficient representation of the time-dependent impact of radiation on tumor cells. (*66, 67*)

Tumor response to radiation is heavily influenced by oxygen saturation (SpO_2_ or %sO_2_), as hypoxic regions are more resistant to radiation-induced damage. SpO_2_ is the fraction of oxygenated hemoglobin relative to total hemoglobin, expressed as a percentage. In healthy systemic circulation, SpO_2_ values typically range from 95–100% (*68*), while in tumors, these values are significantly lower (10–40%) due to poor vascularization, abnormal blood flow, and high metabolic demand for O_2_, leading to hypoxia (*69-71*). AVO is repurposed in this study to reduce tumor oxygen consumption and temporarily increase SpO_2_ levels in hypoxic tumors. This alleviation of hypoxia enhances tumor sensitivity to radiotherapy, forming the basis for our model where AVO pharmacokinetics modulate SpO_2_ dynamically to improve treatment efficacy.

To mathematically capture this modulation, SpO_2_$\left( t \right)$ was modeled as a sigmoidal function of AVO concentration, ensuring that oxygen saturation remains within physiological limits:

$\mathrm{Sp}O_{2}\left( t \right)=\frac{100 \cdot\mathrm{SpO}_{2}^{\mathrm{basal}}}{\mathrm{SpO}_{2}^{\mathrm{basal}}+\left( 100-\mathrm{Sp}O_{2}^{\mathrm{basal}} \right)\cdot e^{-\alpha\cdot\mathrm{AVO}\left( t \right)}}$,	**(S3)**

where, $\mathrm{Sp}O_{2}^{\mathrm{basal}}$ represents the steady-state oxygen saturation in hypoxic tumors, 100% is the physiological maximum oxygen saturation, and $\alpha$ determines how sensitively the oxygen saturation responds to increases in AVO concentration. A higher $\alpha$ value indicates that even modest increases in AVO levels substantially improve oxygenation.

This sigmoidal function ensures that $\mathrm{Sp}O_{2}\left( t \right)$ remains between $\mathrm{SpO}_{2}^{\mathrm{basal}}$ (under hypoxic conditions) and the physiological upper limit of 100%. It provides a smooth transition from hypoxic conditions to enhanced oxygenation as a function of AVO concentration, consistent with physiological constraints.

The systemic pharmacokinetics of AVO following oral administration is described using a one-compartment model. Given the tumor growth timescale of several weeks, absorption dynamics were considered negligible, and the AVO concentration was simplified to a first-order elimination model:

$\mathrm{AVO}\left( t \right)=\sum_{i=1}^{N} \mathrm{Dose}_{\mathrm{AVO}}\cdot\frac{F}{V_{d}}\cdot e^{- \frac{\mathrm{Cl}}{V_{d}} \cdot\left( t-t_{i} \right)}\cdot H\left( t-t_{i} \right)$,	**(S4)**

where $\mathrm{Dose}_{\mathrm{AVO}}$ represents the administered dose of AVO, $F$ is the drug bioavailability via the oral route, $V_{d}$ is the volume of distribution of AVO, $\mathrm{Cl}$ is systemic clearance of AVO, and $t_{i}$ represents the time of the $i$-th dose administration, with $i=1,2,\ldots,N$. The Heaviside function $H\left( t-t_{i} \right)$ ensures the concentration is zero before drug administration. The summation accounts for multiple doses administered over the treatment duration. This approach captures the cumulative effect of repeated dosing on AVO concentration, allowing its systemic concentration to dynamically modulate SpO_2_ within the tumor.

Radiosensitivity is subsequently modulated by $\mathrm{Sp}O_{2}\left( t \right)$, which dynamically adjusts $D_{50}^{\mathrm{effective}}$, the dose of radiation required to achieve half-maximal tumor cell death under current oxygen conditions. This adjustment encapsulates the hypothesis that oxygen enhances the radiosensitivity of tumor cells:

$D_{50}^{\mathrm{effective}}=\frac{D_{50}^{\mathrm{theoretical}}}{\left( \frac{\mathrm{Sp}O_{2}\left( t \right)}{100} \right)^{\beta}}$,	**(S5)**

where, $D_{50}^{\mathrm{theoretical}}$ is the theoretical radiation dose required for half-maximal tumor cell death under fully normoxic conditions ($\mathrm{Sp}O_{2}$=$100\%$) and $\beta$ is a dimensionless parameter that quantifies the sensitivity of $D_{50}^{\mathrm{effective}}$ to deviations in oxygen saturation from normoxia.

Since $\mathrm{Sp}O_{2}\left( t \right)$<100% under hypoxic conditions, the denominator $\left( \frac{\mathrm{Sp}O_{2}\left( t \right)}{100} \right)^{\beta}$ is always less than one, resulting in $D_{50}^{\mathrm{effective}}$>$D_{50}^{\mathrm{theoretical}}$. This indicates that the potency of radiation is reduced under hypoxia compared to normoxia, requiring higher doses of radiation to achieve equivalent therapeutic effects. This relationship emphasizes the critical role of tumor oxygenation in enhancing radiation efficacy and underscores the importance of AVO in alleviating hypoxia.

**Parameter Calibration**

The model was solved numerically as an initial value problem using MATLAB (version R2024a). AVO was administered daily at a dose ($\mathrm{Dose}_{\mathrm{AVO}}$) of 50 mg/kg for one week prior to radiation, with the first dose given on day 1. Radiation was delivered as a single external beam dose on day 8. At the start of the simulation ($t=0$), no radiation had been administered ($\mathrm{Dose}_{\mathrm{rad}}=0$), and a baseline oxygen saturation ($\mathrm{Sp}O_{2}^{\mathrm{basal}}$) of 30% was assumed. Zero initial tumor radiation dose ($\mathrm{Dose}_{\mathrm{rad}}$=0) and a baseline oxygen saturation ($\mathrm{Sp}O_{2}^{\mathrm{basal}}$) of 30% were assumed. The model was solved using the *ode15s* solver, selected for its ability to handle stiff systems often encountered in coupled pharmacokinetic and biological response models.

Four key parameters were fitted based on preclinical data: $\gamma$, $\delta_{\max}$, $\beta$, and $\alpha$. These parameters were optimized using non-linear least squares fitting via MATLAB’s *lsqcurvefit* function, with biologically and physiologically plausible constraints on parameter values. Individual tumor growth curves were fitted for each mouse, using $V_{0}$ as the mouse-specific initial condition. The mean and standard deviation for each parameter were calculated from the resulting distributions, ensuring robust parameter estimates. Parameters not directly fitted were obtained from literature or scaled allometrically between mice and humans. **Table S1** summarizes the parameters, their estimated values, and corresponding sources. This calibration approach integrates experimental data with physiologically informed constraints, enabling accurate modeling of tumor dynamics and treatment efficacy.

**Derivation of Tumor Static Exposure Curve**

The Tumor Static Exposure (TSE) curve quantifies the relationship between the AVO dose ($\mathrm{Dose}_{\mathrm{AVO}}$) and the radiation dose ($\mathrm{Dose}_{\mathrm{rad}}$) required to achieve tumor stasis, defined as a condition where tumor growth is balanced by treatment-induced cell death (${dV\left( t \right)}/{dt=0}$). The derivation builds on the tumor growth model (**Equations S1-S5**) described in the main text.

Under tumor stasis (${dV\left( t \right)}/{dt=0}$), the tumor growth term in Equation S1 balances the radiation-induced tumor cell death term, resulting in:

$\gamma=\delta_{\max}\cdot\frac{D\left( t \right)}{D_{50}^{\mathrm{effective}}+D\left( t \right)}$.

Solving for the required radiation dose ($D\left( t \right)$), we obtain:

$D\left( t \right)=\frac{\gamma}{\delta_{\max}-\gamma}\cdot D_{50}^{\mathrm{effective}}$,

where $D_{50}^{\mathrm{effective}}$, is given in Equation S5:

$D_{50}^{\mathrm{effective}}=\frac{D_{50}^{\mathrm{theoretical}}}{\left( \frac{SpO2(t)}{100} \right)^{\beta}}$.

In Equation S2, $D\left( t \right)$ describes the time-dependent decay of radiation dose due to biological processes like DNA repair. For the TSE derivation, we replace $D\left( t \right)$ with $\mathrm{Dose}_{\mathrm{rad}}$, which is the administered dose of radiation. This substitution focuses on the initial effect of radiation and aligns with experimental and clinical frameworks, where total administered dose is the primary variable. Thus, by replacing $D\left( t \right)$ with $\mathrm{Dose}_{\mathrm{rad}}$ and substituting the expression for $D_{50}^{\mathrm{effective}}$, we arrive at the following relation:

$\mathrm{Dose}_{\mathrm{rad}}=\frac{\gamma}{\delta_{\max}-\gamma}\cdot\frac{D_{50}^{\mathrm{theoretical}}}{\left( \frac{SpO2(t)}{100} \right)^{\beta}}$.

Tumor oxygen saturation ($SpO2(t)$) dynamically improves with AVO administration, as described in Equation S3. As a simplification, we evaluate the pharmacodynamic effects of AVO using a single dose and assume that the peak concentration ($\mathrm{AVO}_{\mathrm{peak}}$), occurring immediately after administration due to the assumption of instantaneous absorption, captures its effects on oxygen modulation and subsequent tumor radiosensitivity. The peak AVO concentration ($\mathrm{AVO}_{\mathrm{peak}}$) is:

$\mathrm{AVO}_{\mathrm{peak}}=\frac{\mathrm{Dose}_{\mathrm{AVO}}\cdot F}{V_{d}}$.

Substituting $\mathrm{AVO}_{\mathrm{peak}}$ into Equation S3 gives:

$\mathrm{Sp}O_{2}\left( t \right)=\frac{100 \cdot\mathrm{SpO}_{2}^{\mathrm{basal}}}{\mathrm{SpO}_{2}^{\mathrm{basal}}+\left( 100-\mathrm{Sp}O_{2}^{\mathrm{basal}} \right)\cdot e^{-\alpha\cdot\frac{\mathrm{Dose}_{\mathrm{AVO}}\cdot F}{V_{d}}}}$.

Substituting $\mathrm{Sp}O_{2}\left( t \right)$ into the expression for $\mathrm{Dose}_{\mathrm{rad}}$ gives the final TSE curve:

$\mathrm{Dose}_{\mathrm{rad}}=\frac{\gamma}{\delta_{\max}-\gamma}\cdot D_{50}^{\mathrm{theoretical}}{\cdot\left( \frac{\mathrm{Sp}O_{2}^{\mathrm{basal}}+\left( 100-\mathrm{Sp}O_{2}^{\mathrm{basal}} \right)\cdot e^{-\alpha\cdot\frac{\mathrm{Dose}_{\mathrm{AVO}}\cdot F}{V_{d}}}}{\mathrm{Sp}O_{2}^{\mathrm{basal}}} \right)}^{\beta}$.

This expression quantifies the interplay between radiation dose and AVO dose, assuming that AVO’s immediate effects on tumor oxygenation are described by its peak concentration following a single dose. By simplifying the radiation dynamics to focus on cumulative dose ($\mathrm{Dose}_{\mathrm{rad}}$), the model aligns with experimental and clinical frameworks, where total dose rather than time-dependent decay is the primary variable of interest.

***Optimization of Radiation Start Time and AVO Dosing Frequency.*** To assess the impact of radiation timing, simulations were performed for radiation initiation days ranging from day 0 (simultaneous with AVO start) to day 30 (radiation starting 30 days after AVO initiation). AVO was administered daily at a dose of 10 mg/kg for four weeks, while radiation was delivered at 2 Gy per fraction, five days per week, for four weeks. In addition to daily administration, an alternate dosing schedule of AVO administered once every two days (10 mg/kg) was also tested to assess the impact of administration frequency on timing optimization. For each start day scenario, $\Delta d$ was calculated for all 500 virtual patients one month after treatment completion, and the fraction of patients achieving PR was determined. The optimal start time was defined as the day that maximized the fraction of PR across the population.

***Optimization of Combination Doses.*** To identify clinically optimal dose combinations of AVO and radiation, simulations were conducted for AVO doses ranging from 0.1–100 mg/kg and total radiation doses ranging from 0.1–100 Gy. AVO was administered daily for four weeks. Radiation was delivered over four weeks (five days per week, i.e., 20 fractions), with the dose per fraction adjusted for each combination to achieve the specified total radiation dose. For example, a total radiation dose of 40 Gy was divided into 20 fractions of 2 Gy each, while a total dose of 80 Gy was divided into 20 fractions of 4 Gy each.

A total of N=2,500 dose combinations were simulated, with 50 evenly spaced values chosen for both AVO doses and total radiation doses. For each dose combination, the tumor response was simulated across the virtual patient population, and the fraction of patients achieving a PR was calculated one month after the end of treatment. The optimization goal was to identify dose combinations that minimized total radiation utilization while maintaining high tumor control in the presence of AVO. The results were visualized as a contour plot, with color grading representing the fraction of PR for each combination.

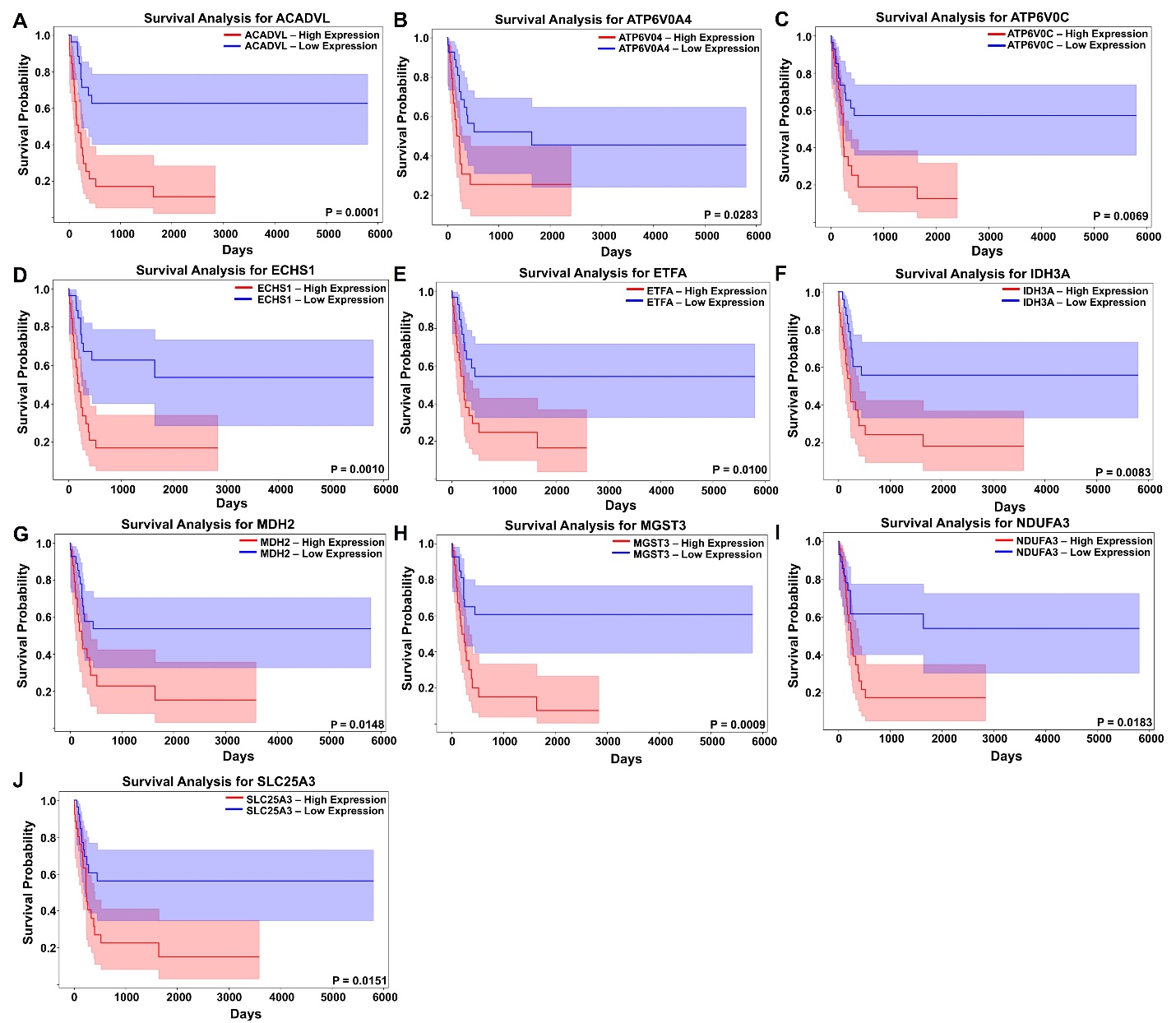

**Fig. S1.** Kaplan-Meier survival analysis showed that patients with low ACADVL, ATP6VOA4, ATP6V0C, ECHS1, ETFA, IDH3A, MDH2, MGST3, NDUFA3 and SLC25A3 expression had a significantly better survival rate compared to those with high expression.

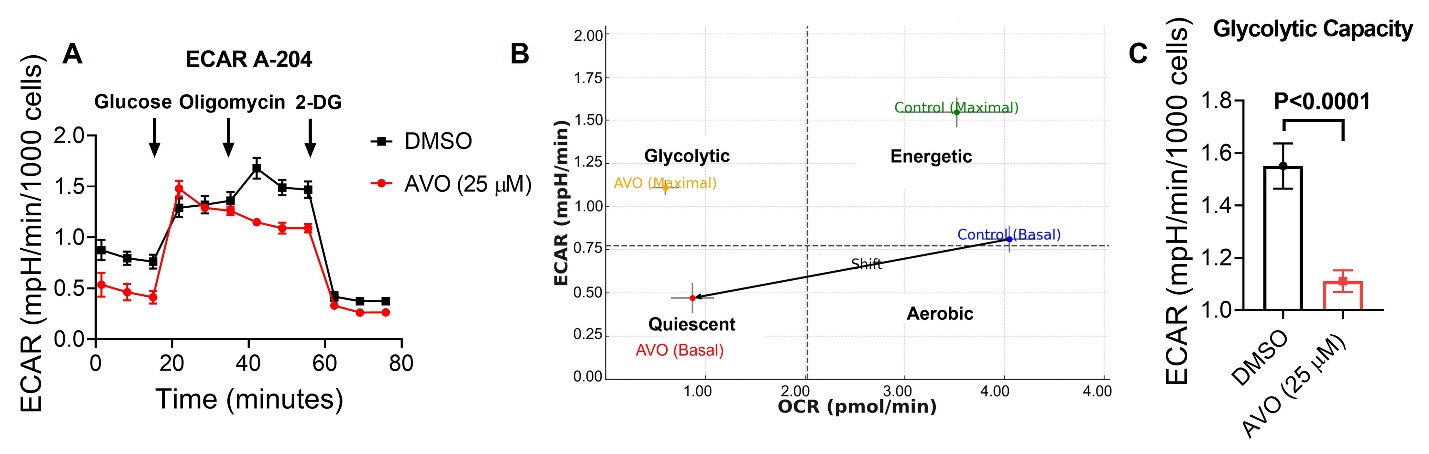

**Fig. S2.** AVO treatment does not show a significant glycolytic shift in A-204 cells. **(A)** ECAR traces of A-204 cells treated with vehicle (DMSO) or AVO. **(B)** ECAR and OCR were plotted on the same graph. Treatment with AVO (25 µM) transitions A-204 cells from an aerobic/energetic state to a quiescent state. **(C)** Comparison of glycolytic capacity in A-204 cells treated with DMSO or AVO. Data presented as mean ± S.D.

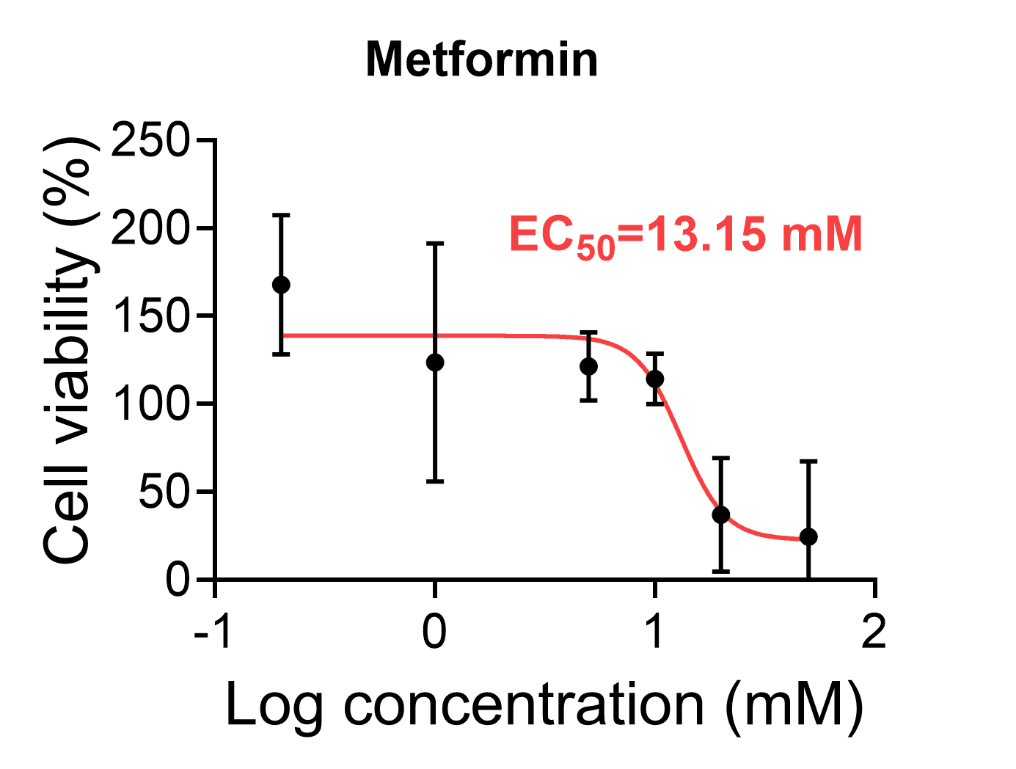

**Fig. S3.** Dose response curves depict the effect of Metformin on viability of A-204 cells as tested by MTT assay. Data presented as mean ± S.D.

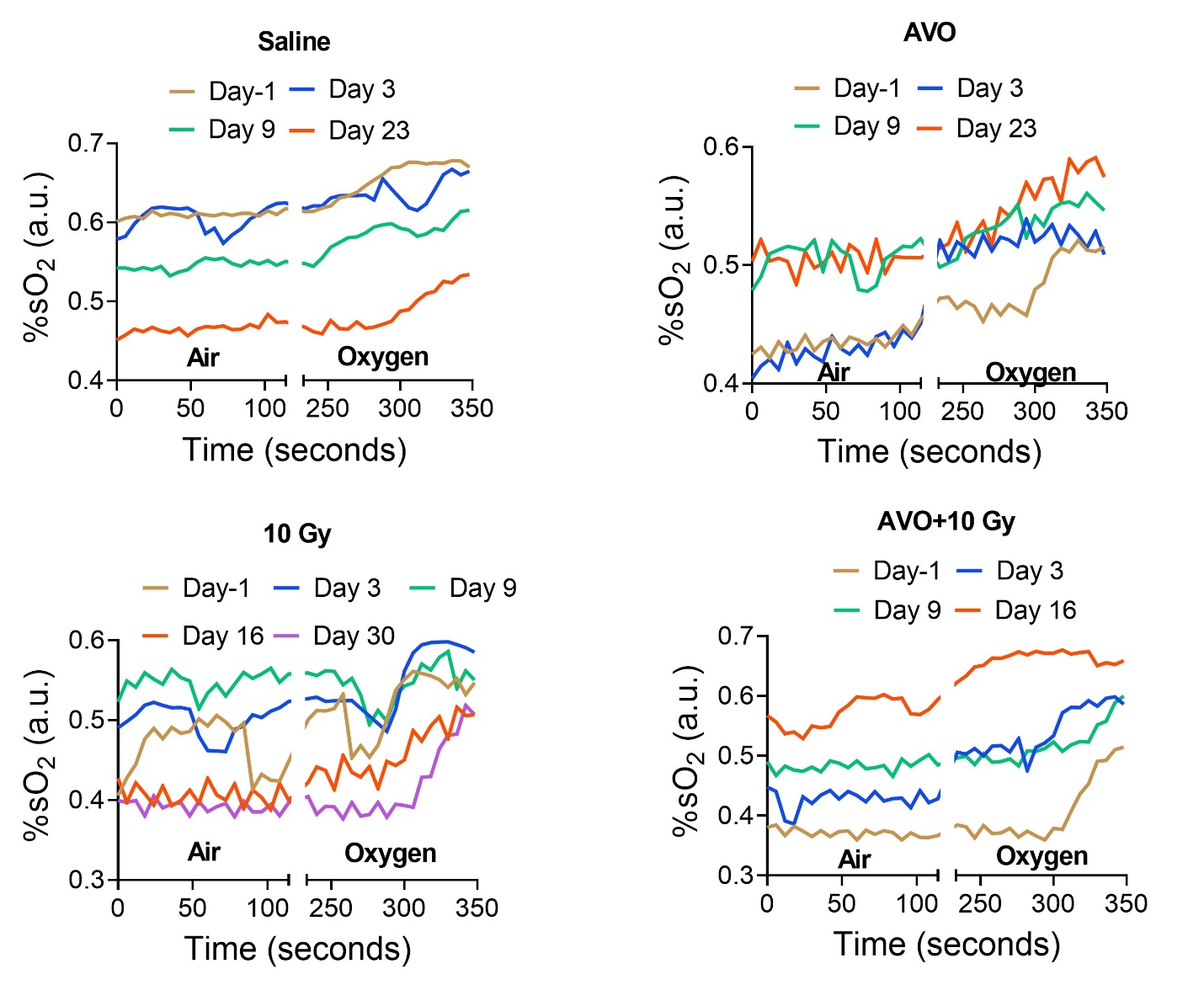

**Fig. S4**. Curves depicting changes in oxygen saturation in response to oxygen challenge during OE-MSOT.

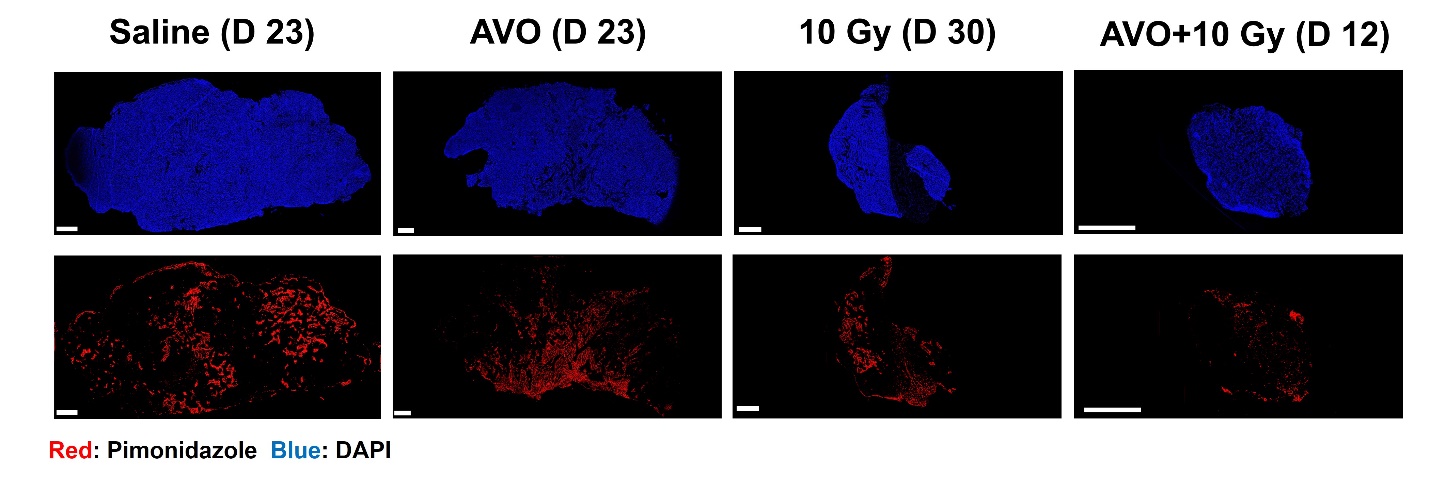

**Fig. S5**. Whole tissue immunofluorescence imaging depicting pimonidazole staining of tumors from different groups. Scale bar = 1000 µm.

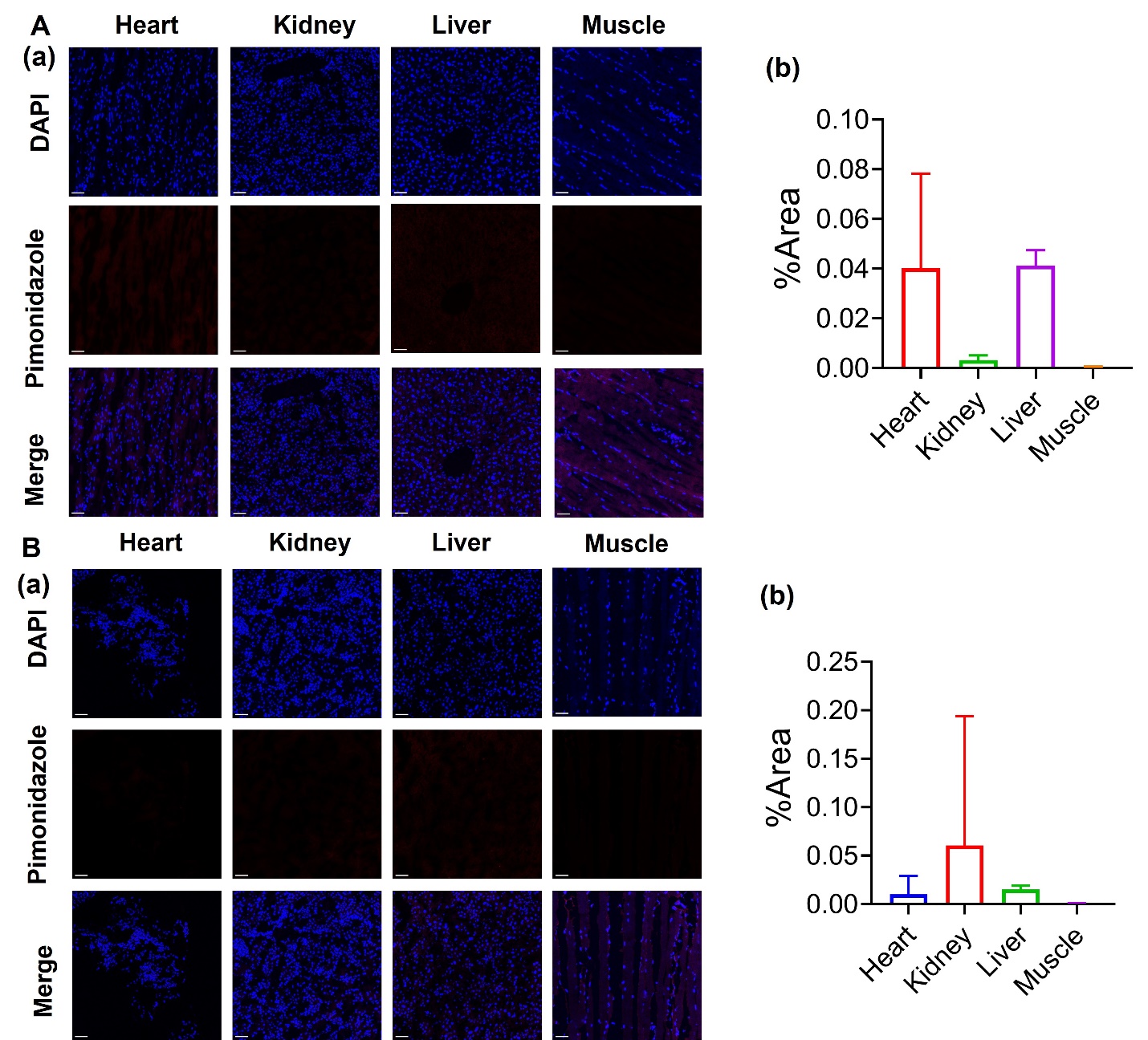

**Fig. S6.** The confocal images (a) and quantitative analysis (b) of pimonidazole staining of heart, kidney, liver, and muscle tissue of (A) saline-treated and (B) AVO+10 Gy-treated group. Scale bar=50 µm. Data presented as mean ± S.D.

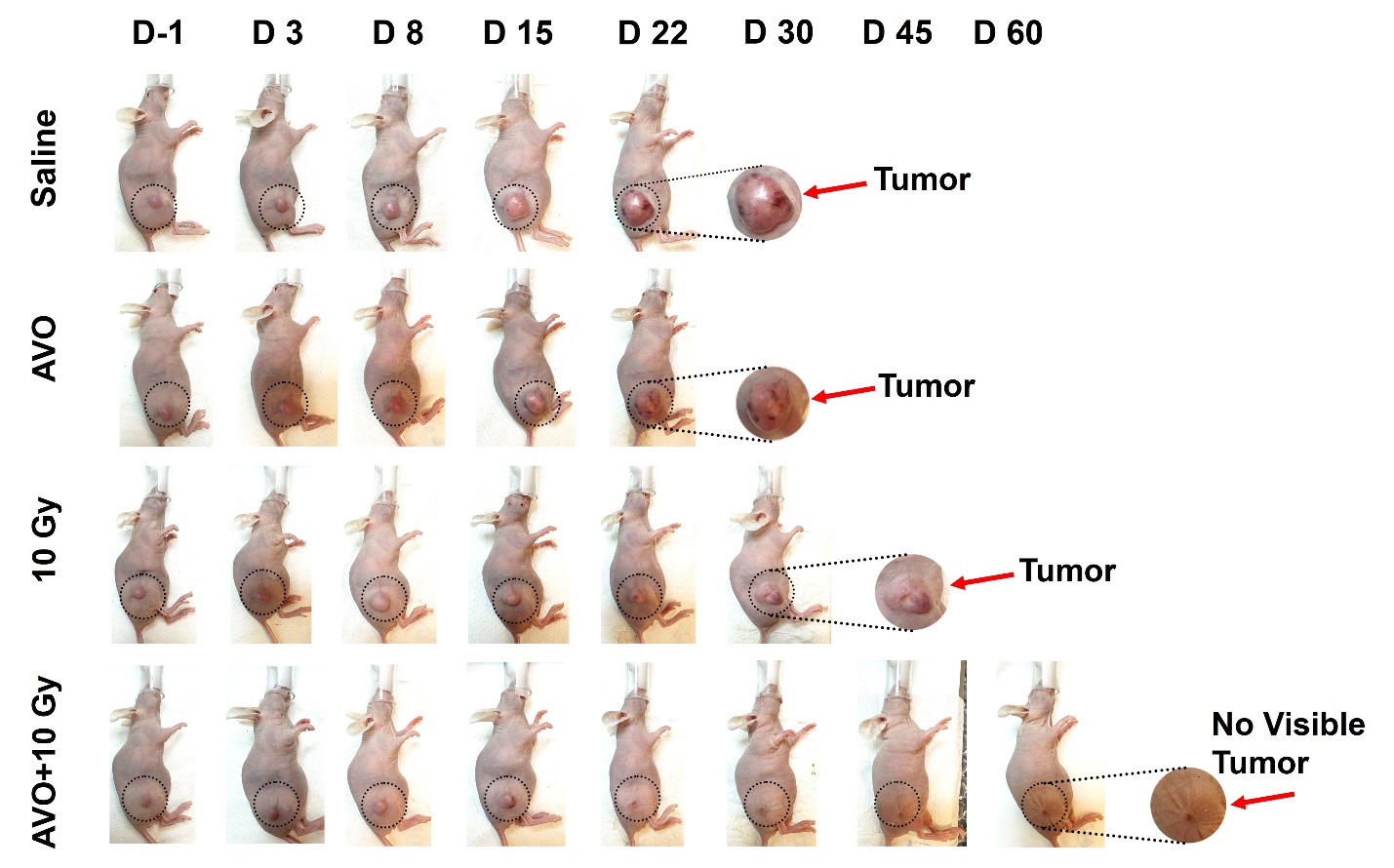

**Fig. S7.** Photographs of A-204 tumor-bearing mice taken at various timepoints during the course of different treatments. In the AVO+10 Gy treatment group, the tumor was diminished after Day 16 and only a scar was visible by endpoint.

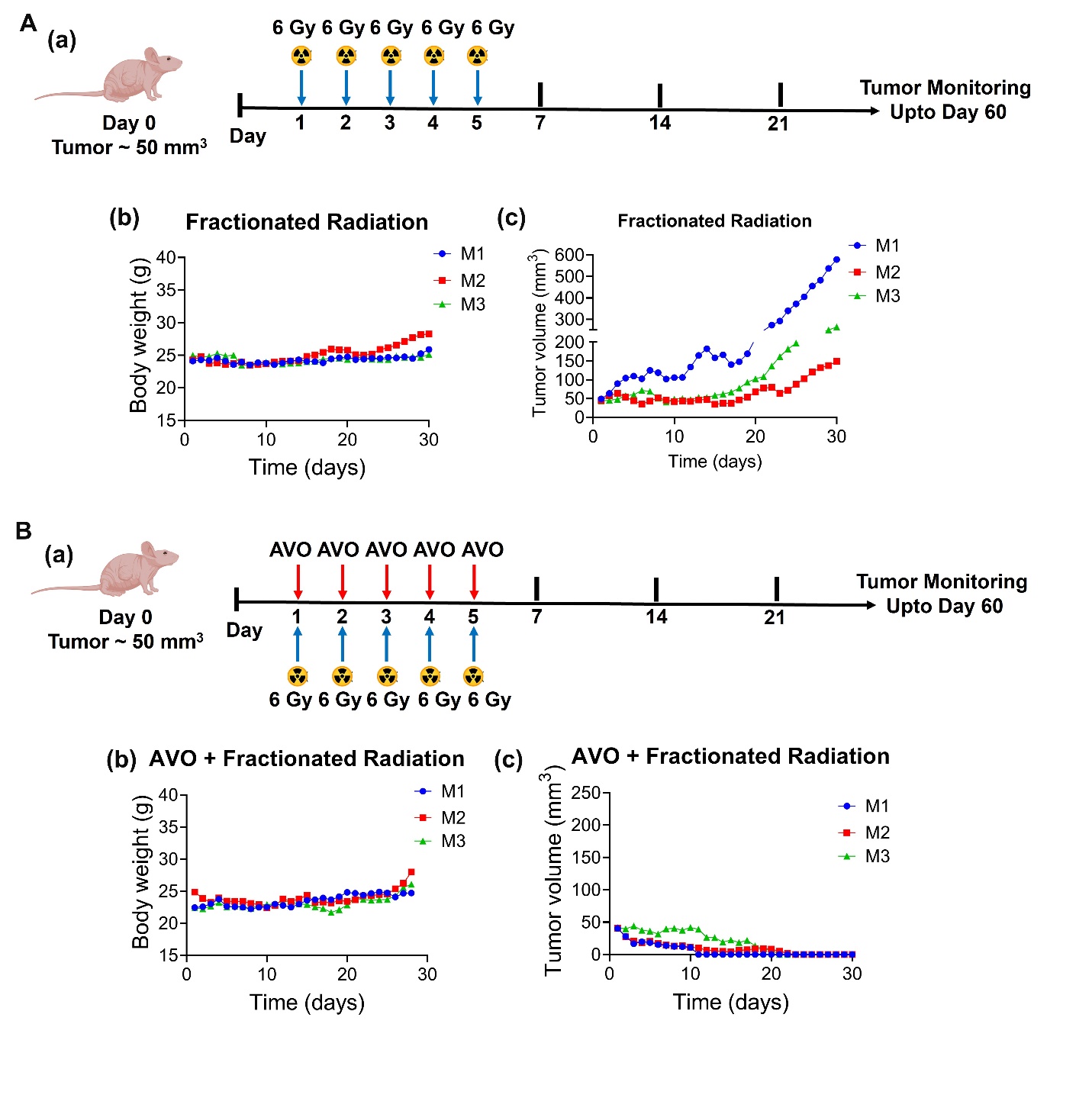

**Fig. S8.** **(A)** (a) Schematic depicting fractionation regimen (5 x 6 Gy dose) of high dose RT administered to A-204 xenografts; (b) body weights changes; and (c) tumor volume changes of individual mice. **(B)** (a) Schematic depicting treatment regimen; (b) body weight changes and (c) tumor volume changes of mice administered with AVO + fractioned radiation treatment groups. (n=3).

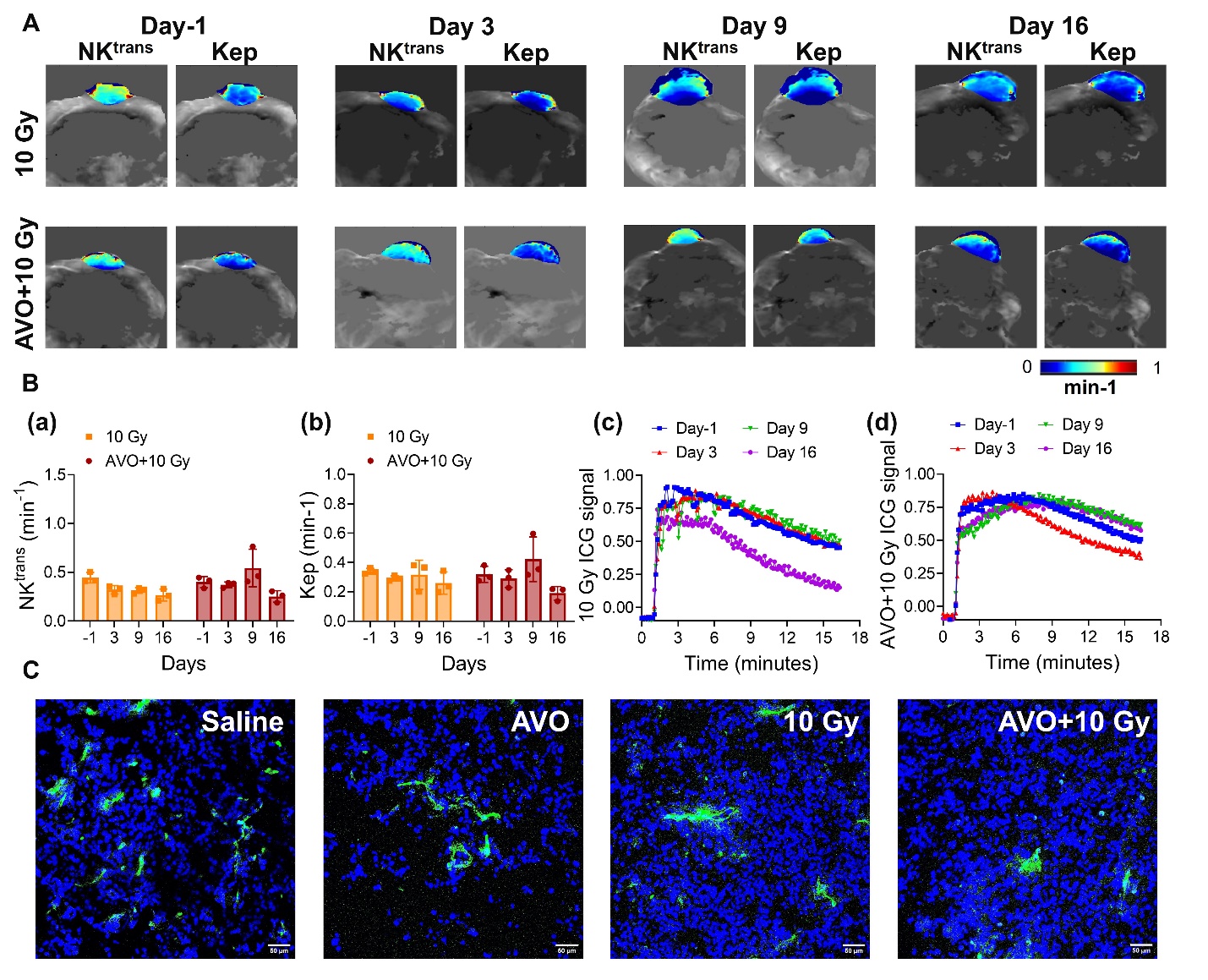

**Fig. S9**. **DCE-MSOT shows that there was no difference in tumor perfusion between 10 Gy and AVO+10 Gy treatment groups**. **(A)** Parametric maps and **(B)** quantification analysis of NK^trans^ and Kep for (a)10 Gy and (b) AVO+10 Gy treatment groups on Day-1, Day 3, Day 9 and Day 16. Kinetic curves depicting changes in ICG signals in tumors treated with (c) 10 Gy or (d) AVO+10 Gy. **(C)** Confocal microscopy of CD31 stained tumor sections excised at Day 9 from different treatment groups. Scale bar=50 µm.

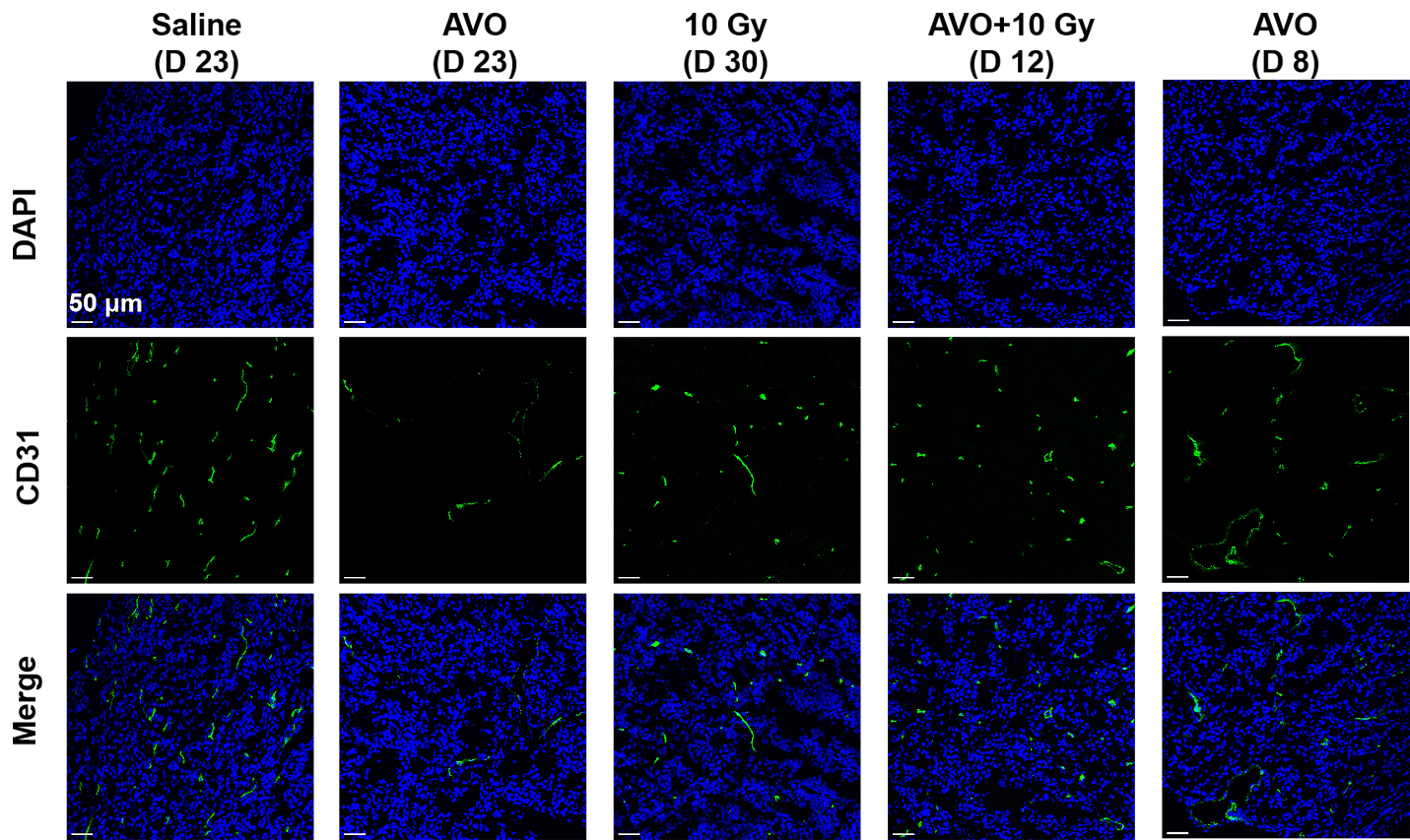

**Fig. S10.** Confocal microscopy images of CD31 staining of tumor sections excised from mid- or post-treatment timepoints. Scale bar=50 µm.

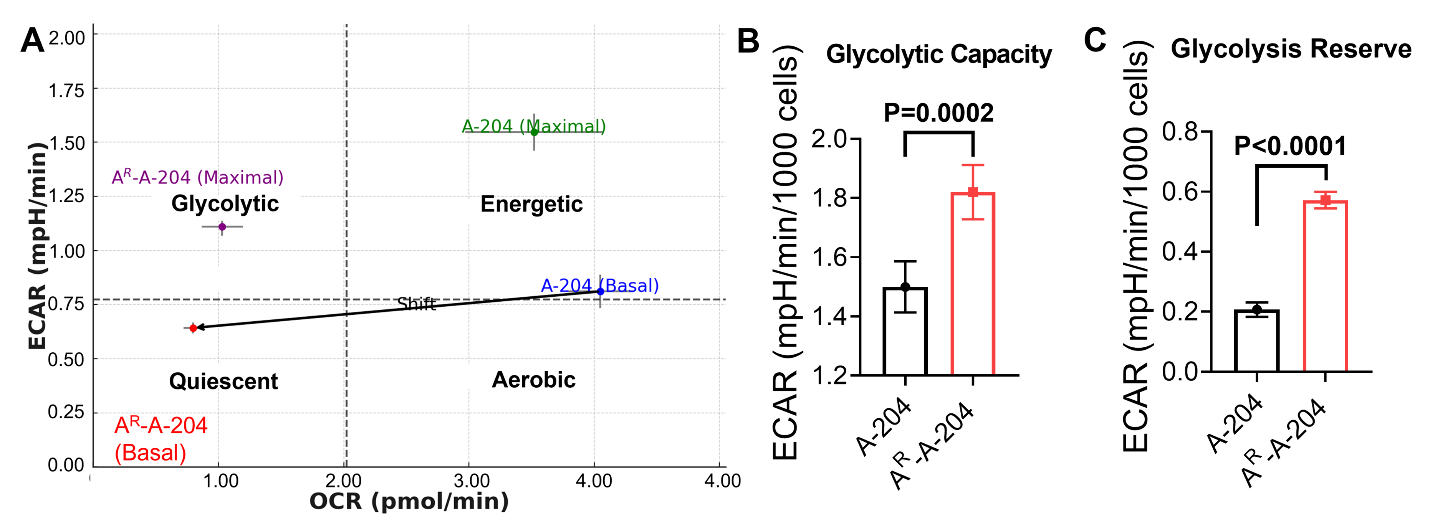

**Fig. S11**. AVO treatment does not show a significant glycolytic shift in A^R^-A-204 cells. **(A)** ECAR and OCR were plotted on the same graph. Following prolonged AVO exposure, A-204 cells shifted from an energetic state to a quiescent state. Comparison of glycolytic capacity **(B)** and glycolysis reserve **(C)** between A-204 and A^R^-A-204 cells. Data presented as mean ± S.D.

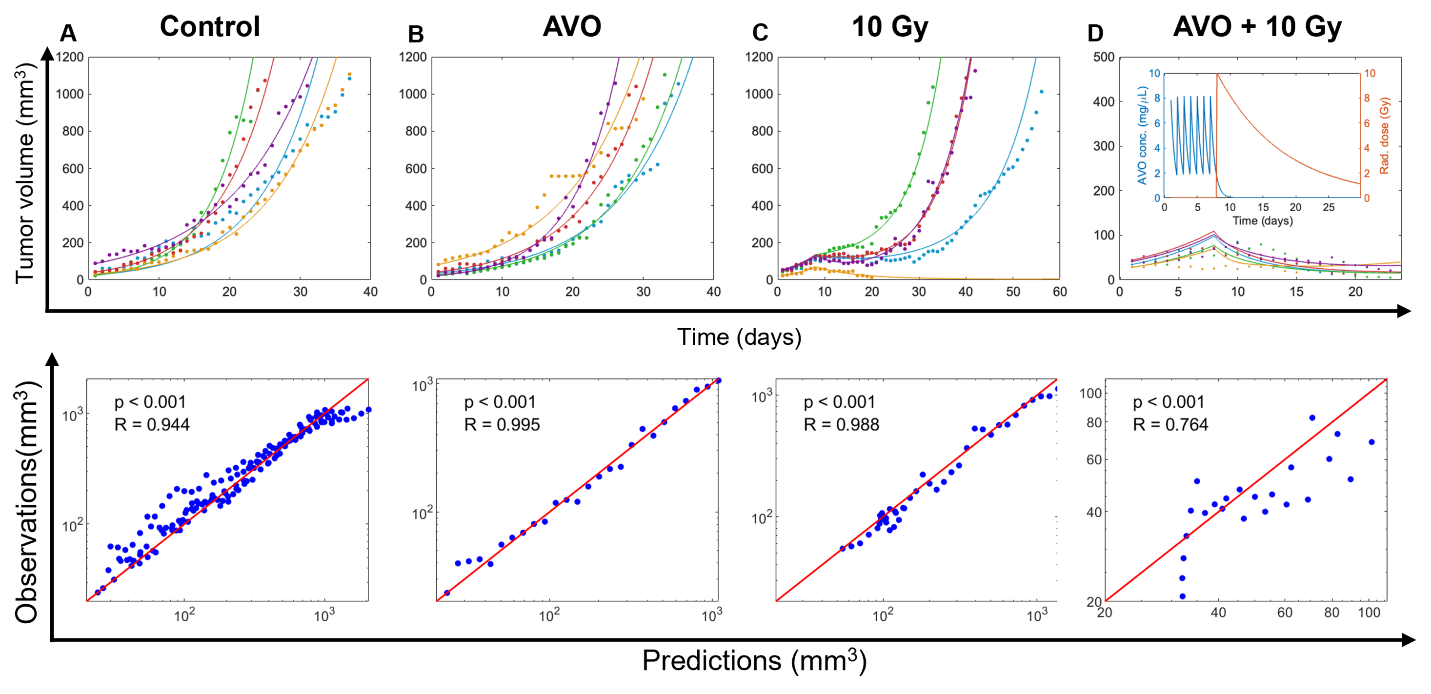

**Fig. S12**. **(A–D; Upper Panel)** Model fits to mouse tumor growth kinetics under different therapeutic regimens: **(A)** control, **(B)** AVO monotherapy (50 mg/kg/day for 7 days), **(C)** RT monotherapy (10 Gy single dose on day 8), and **(D)** combination therapy (AVO + RT) following the same regimen as the monotherapies. The inset in **(D)** shows the modeled AVO concentration kinetics (left y-axis) and the modeled biological dissipation of a single radiation dose (right y-axis). (Lower Panel): Pearson correlation analysis comparing predicted and observed tumor volumes under each treatment condition corresponding to figures A-D in the upper panel. Each panel shows the line of identity (red) and individual data points (blue), with associated p-values and correlation coefficients (R) indicating the model’s predictive accuracy.

**Supplementary Tables**

**Table S1. List of model parameters.**

| **Parameter** | **Description** | **Units** | **Value (Mice)** | | | | **Value (Humans)** | **Reference** |
| --- | --- | --- | --- | --- | --- | --- | --- | --- |
|  |  |  | **Control** | **AVO** | **Rad** | **AVO + Rad** |  |  |
| $\gamma$ | Tumor growth rate | d^-1^ | 0.123$\pm$  0.03^***^ | 0.112$\pm$  0.02^***^ | 0.145$\pm$  0.01^***^ | 0.137$\pm$  0.009^***^ | 0.0169 – 0.02^**^ |  |
| $\delta_{\max}$ | Maximum radiation-induced cell death rate | d^-1^ | – | – | 0.518$\pm$  0.21^***^ | 0.518$\pm$  0.04^***^ | 0.057 – 0.07^**^ |  |
| $\alpha$ | Sensitivity factor governing how changes in AVO concentration affect oxygen saturation | L mg^-1^ | – | – | – | 0.0025$\pm$  0.003^***^ | 0.0033 – 5^**^ |  |
| $\beta$ | Exponent determining how changes in oxygen saturation influence radiation potency | – | – | – | 0.88$\pm$  1^***^ | 1.56$\pm$  0.33^***^ | 1.4 – 2.1^**^ |  |
| $\mathrm{SpO}_{2}^{\mathrm{basal}}$ | Steady-state oxygen saturation in tumor | % | 30 | | | | 10 – 40 | (69-71) |
| $\mathrm{Cl}$ | Clearance of AVO | L d^-1^ | 0.0026 – 0.004^*^ | | | | 1.2 – 1.8 | (72) |
| $V_{d}$ | Volume of distribution of AVO | L kg^-1^ | 0.0016^*^ | | | | 5.8 | (72) |
| $F$ | Oral bioavailability of AVO | – | 0.21 | | | | 0.21 – 0.35 | (72, 73) |
| $D_{50}^{\mathrm{theoretical}}$ | Radiation dose for half-maximal death under normoxia | Gy | 1 | | | | 1 | (74) |
| $k$ | Biological decay rate of radiation efficacy | d^-1^ | 0.1 | | | | 0.1 | (74) |

*These parameters were allometrically scaled from humans to mice.

**These parameters were allometrically scaled from mice to humans.
***These parameters were fitted.
